# Alphavirus-induced transcriptional and translational shutoffs play major roles in blocking the formation of stress granules

**DOI:** 10.1101/2023.07.05.547824

**Authors:** Oksana Palchevska, Francisco Dominguez, Elena I. Frolova, Ilya Frolov

## Abstract

Alphavirus infections cause multiple alterations in the intracellular environment that can have both positive and negative effects on viral replication. The Old World alphaviruses, such as Sindbis (SINV), chikungunya (CHIKV), and Semliki Forest viruses, hinder the ability of vertebrate cells to form stress granules (SGs). Previously, this inhibitory function was attributed to the hypervariable domain (HVD) of nsP3, which sequesters the key components of SGs, G3BP1 and G3BP2, and to the nsP3 macro domain. The macro domain possesses ADP-ribosylhydrolase activity, which can diminish the ADP-ribosylation of G3BP1 during viral replication. However, our recent findings do not support the prevailing notions. We demonstrate that the interactions between SINV– or CHIKV-specific nsP3s and G3BPs, and the ADP-ribosylhydrolase activity are not major contributors to the inhibitory process, at least when nsP3 is expressed at biologically relevant levels. Instead, the primary factors responsible for suppressing SG formation are virus-induced transcriptional and translational shutoffs that rapidly develop within the first few hours post infection. Poorly replicating SINV variants carrying mutated nsP3 HVD still inhibit SG development even in the presence of NaAs. Conversely, SINV mutants lacking transcription and/or translation inhibitory functions lose their ability to inhibit SGs, despite expressing high levels of wt nsP3. Moreover, we found that stable cell lines expressing GFP-nsP3 fusions retain the capacity to form SGs when exposed to sodium arsenite. However, our results do not rule out a possibility that additional virus-induced changes in cell biology may contribute to the suppression of SG formation.

**Importance:** Our study highlights the mechanisms behind the cell’s resistance to SG formation after infection with Old World alphaviruses. Shortly after infection, the replication of these viruses hinders the cell’s ability to form SGs, even when exposed to chemical inducers such as sodium arsenite. This resistance is primarily attributed to virus-induced transcriptional and translational shutoffs, rather than interactions between the viral nsP3 and the key components of SGs, G3BP1/2, or the ADP-ribosylhydrolase activity of nsP3 macro domain. While interactions between G3BP and nsP3 are essential for the formation of viral replication complexes, their role in regulating SG development appears to be minimal, if any. Cells harboring replicating virus-specific RNA with modified abilities to inhibit transcription and/or translation, but encoding wt nsP3, retain the capacity for SG development. Understanding these mechanisms of regulation of SG development contributes to our knowledge of viral replication and the intricate relationships between alphaviruses and host cells.

## Introduction

The *Alphavirus* genus in the *Togaviridae* family comprises over 30 known members that are distributed worldwide (1). Most alphaviruses are transmitted by mosquito vectors between vertebrate hosts, causing a variety of human diseases (2). Based on the areas of geographical circulation, alphaviruses are classified into the New World (NW) and the Old World (OW) species. Many NW alphaviruses are encephalitogenic, and their infections in humans may lead to either a lethal outcome or neurological sequelae (3). The diseases caused by the OW alphaviruses are characterized by the development of polyarthritis that can continue for months (4). Despite the unquestionable public health threat, the basis of alphavirus pathogenesis at the molecular, cellular, and organismal levels remains insufficiently understood.

Alphavirus genomic RNA (G RNA) encodes a handful of proteins. Four viral nonstructural proteins (nsP1-to-4) are translated directly from the G RNA as polyproteins P123 and P1234 (1). Their sequential processing by nsP2-associated protease activity regulates the functions of viral replication complexes (vRCs) during negative and positive strand RNA syntheses (5–8). Early after infection, the vRCs contain partially processed nsPs (P123+nsP4) and synthesize the negative strand of the viral genome to form double-stranded RNA (dsRNA) replication intermediates. Later, fully processed nsPs within the vRCs produce viral G RNAs and subgenomic (SG) RNAs. The latter RNA encodes a precursor polyprotein for viral structural proteins (9, 10).

The synthesis of alphavirus structural and nonstructural proteins, as well as RNA replication induce multiple changes in cellular biology. These modifications create an environment that facilitates viral replication and hampers the activation of the antiviral response. Shortly after infection, both the OW and NW alphaviruses induce transcriptional shutoff (11–15), which is a critical mechanism for inhibiting the expression of cellular genes involved in antiviral and innate immune responses. Some of the OW alphaviruses, such as Sindbis (SINV) and Semliki Forest (SFV) viruses, also exhibit highly efficient translation inhibition (16, 17), which additionally suppresses the antiviral response.

Like many other viruses, alphaviruses generate substantial amounts of dsRNAs (18, 19), which can act as one of the pathogen-associated molecular patterns (PAMPs). The isolation of these dsRNAs into membranous spherules (19, 20) formed during viral RNA replication appears to be incomplete and this results in the activation of cellular pattern recognition receptors (PRRs), such as RIG-I, MDA-5, and PKR (17, 21, 22). Interaction of PKR with dsRNA leads to its phosphorylation, dimerization, and subsequent phosphorylation of the translation initiation factor eIF-2α. This PKR-induced phosphorylation negatively affects the availability of the non-phosphorylated form of eIF-2α, which is essential for the assembly of the ternary eIF2/tRNA_i_^Met^/GTP complexes. Ultimately, a large portion of cellular mRNAs becomes stalled in 48S initiation complexes, impairing the translation of host proteins. Phosphorylation of eIF-2α can be also mediated by PKR-like endoplasmic reticulum kinase (PERK), heme-regulated inhibitor kinase (HRI) and general control non-repressed 2 kinase (GCN2) (23). During viral replication, the PKR-dependent mechanism is likely the most important contributor to the inhibition of cellular translation. However, in SINV-infected cells, the second component of translation inhibition is PKR-independent (21) and remains poorly understood. Interestingly, the translation of SINV SG RNA, which contains a translational enhancer in its 5’ UTR, remains efficient despite the overall translation inhibition.

The accumulated stalled 48S initiation complexes attract large amounts of cellular RNA-binding proteins, leading to the formation of stress granules (SGs) (24, 25). These dynamic membraneless organelles serve as sites for the accumulation of mRNPs and stalled translation initiation complexes during translational arrests induced by a variety of stimuli. SGs have the potential to interfere with viral replication by sequestering virus-specific mRNAs. However, some viruses, including alphaviruses, have developed specific strategies to counteract SG assembly (26, 27). The OW alphaviruses, such as SFV and chikungunya virus (CHIKV), transiently induce SG formation at early times post infection (p.i.), but subsequently inhibit SG assembly even in response to external inducers at later times (28, 29). Two mechanisms of the inhibition of SG assembly have been proposed. The first hypothesis is based on the ability of the hypervariable domain (HVD) of the nonstructural protein nsP3 to bind the main SG components, Ras-GTPase activating SH3 domain-binding proteins (G3BP1 and G3BP2) (20, 30–33). The nsP3-G3BP complexes were proposed to sequester the entire pool of cellular G3BPs, thereby altering SG development and indirectly promoting viral replication (32, 34). Another recent study suggests an alternative hypothesis that the macro domain of alphavirus nsP3 downregulates SG formation in CHIKV-infected cells by reducing ADP-ribosylation of G3BP1 (29).

In our study, we focused on investigating SG assembly in SINV– and CHIKV-infected cells. Our data demonstrate that alterations in SINV nsP3 interactions with G3BPs significantly impair SINV replication. However, the lack of binding between G3BPs and nsP3 HVD does not restore the cells’ ability to form SGs during replication of SINV mutants. Even under chemically induced stress, cells infected with SINV nsP3 HVD mutants remain unable to assemble SGs. Furthermore, we show that the expression of SINV nsP3 alone does not prevent SG assembly in response to oxidative stress inducer, sodium arsenite (NaAs). Thus, the sequestration of G3BPs into nsP3-containing protein complexes and the ADP-ribosylhydrolase activity of the macro domain are not the primary contributors to the inhibition of SG assembly. Instead, virus-induced transcriptional and/or translational inhibition plays critical roles in preventing SG formation during viral replication and NaAs treatment.

## Results

### SINV and CHIKV infections hinder the assembly of SGs

Previous studies have suggested that SFV infection efficiently triggers SG formation at the early times p.i., and they gradually disassemble as the infection progresses (27). Therefore, our initial experiments aimed to compare SG formation during the replication of two other OW alphaviruses, SINV and CHIKV, in NIH 3T3 cells that are highly permissive for alphavirus infections.

At 3 h p.i. with SINV/GFP, only approximately 3% of cells exhibited the presence of SGs. Additionally, as depicted in the representative image in Fig. 1, the infected GFP-positive cells displayed a lower number of SGs per cell compared to those mock-infected and treated with NaAs. No SGs were detected in CHIKV-infected cells at 3 h p.i. (Fig. 2), and we did not observe any SGs at 6 h p.i. with either virus.

**Fig. 1.**
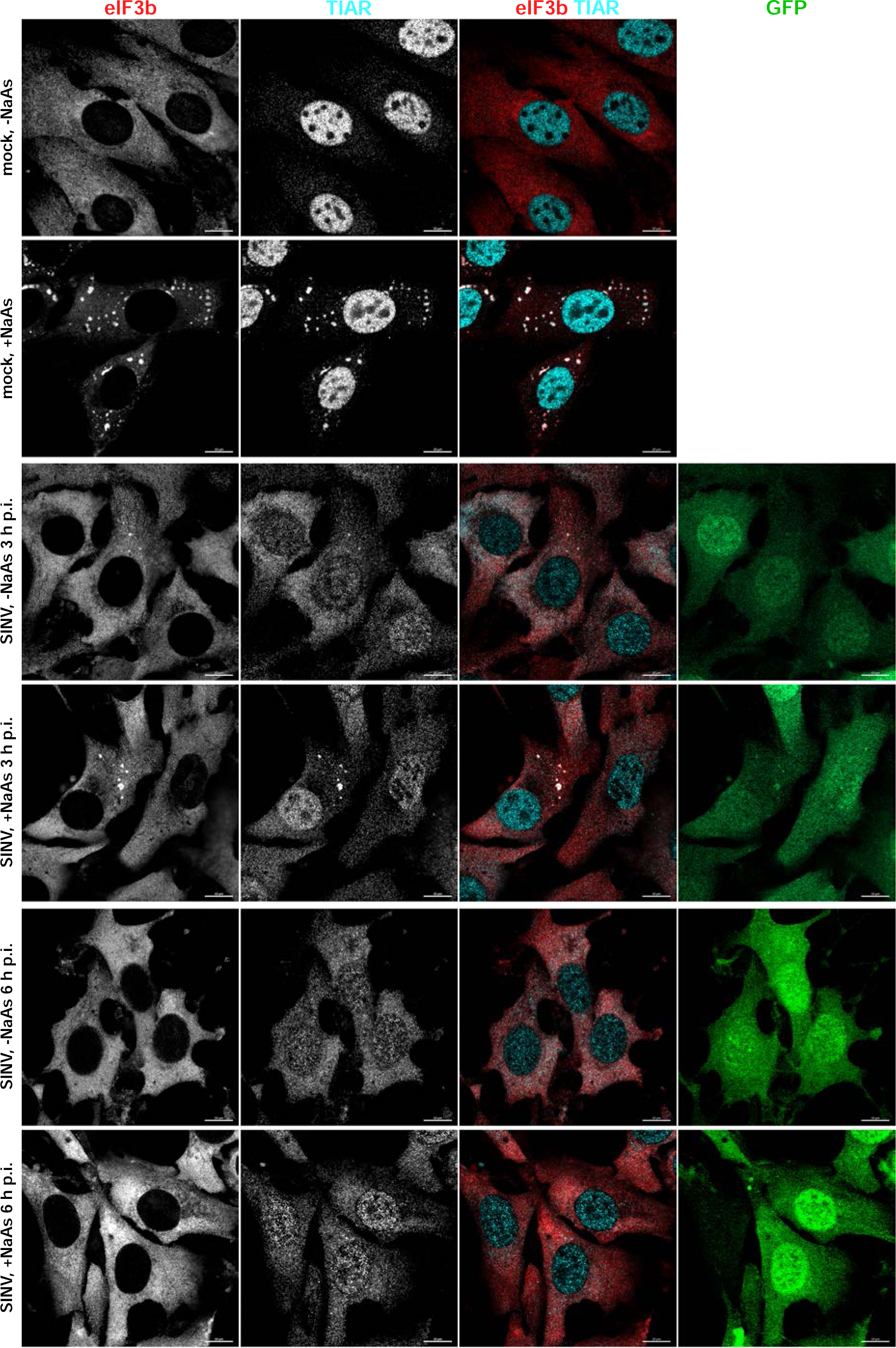
Replication of SINV rapidly makes cells resistant to SG formation. NIH 3T3 cells were infected with SINV/GFP at an MOI of 10 PFU/cell and then, at the indicated times p.i., were either mock-treated or exposed to NaAs as described in Materials and Methods. Cells were immunostained with antibodies specific to SG components. Bars correspond to 10 μM. The experiment was reproducibly repeated several times.

**Fig. 2.**
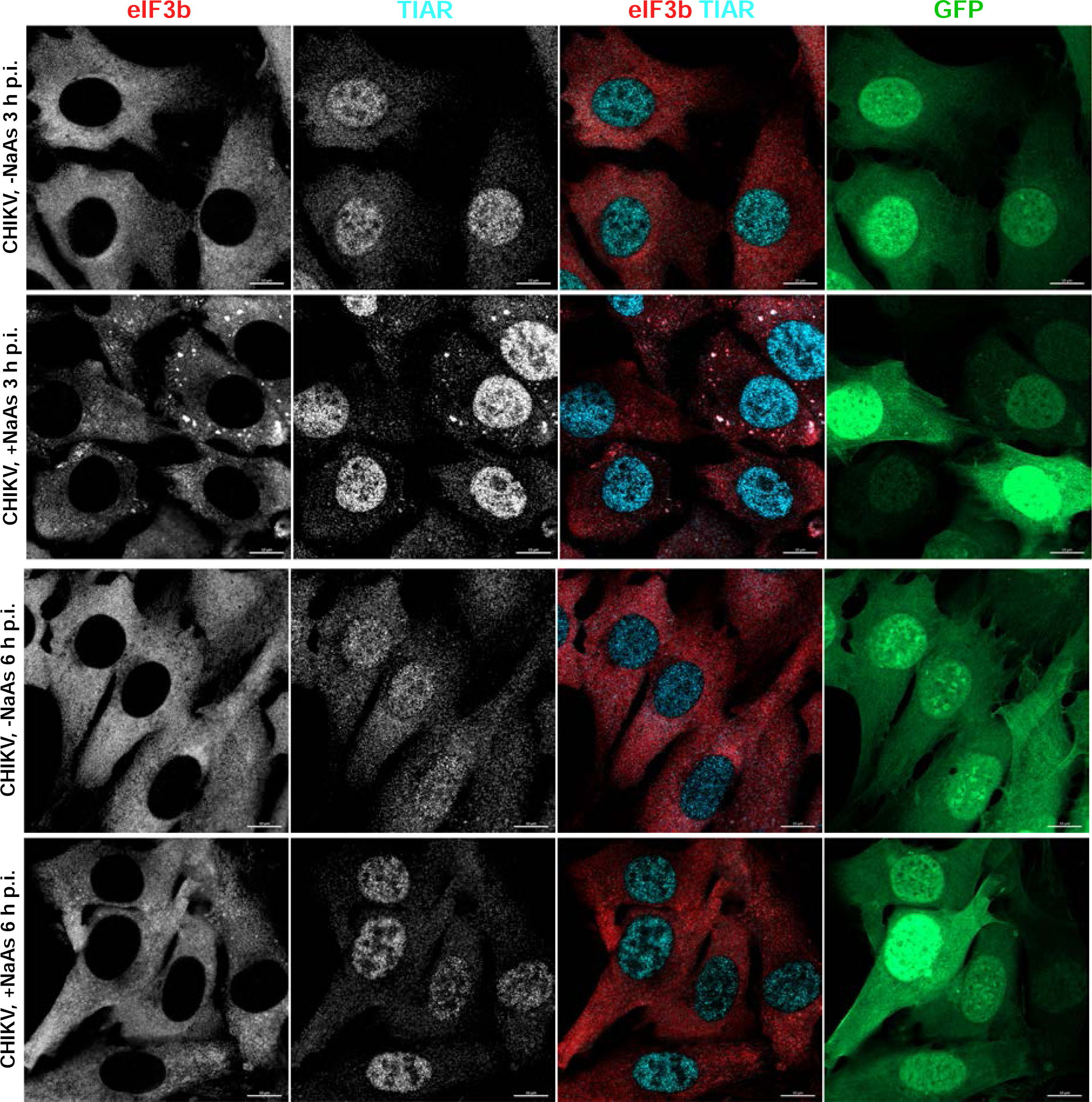
CHIKV infection also makes cells resistant to SG formation. NIH 3T3 cells were infected with CHIKV/GFP at an MOI of 10 PFU/cell. At the indicated times p.i., they were exposed to NaAs. Cells were immunostained with the indicated antibodies specific to SG components. Bars correspond to 10 μM. The experiment was reproducibly repeated multiple times.

The formation of SGs requires the phosphorylation of eIF2α. Western blot (WB) analysis of eIF2α phosphorylation revealed a rapid accumulation of the phosphorylated form of eIF2α (p-eIF2α) in SINV-infected cells (Fig. 3). By 6 h p.i., the level of p-eIF2α was similar to that found in cells treated with NaAs. Surprisingly, CHIKV infection induced less efficient eIF2α phosphorylation, which may explain the absence of SGs in CHIKV-infected cells at 3 and 6 h p.i. However, both SINV and CHIKV infections actively inhibited SG formation upon treatment of the cells with NaAs (Figs. 1 and 2). At 3 h p.i., only approximately 15% of SINV-infected cells and around 70% of CHIKV-infected cells formed SGs in response to NaAs treatment. At 6 h p.i., NaAs failed to induce SGs in both SINV– and CHIKV-infected cells (Figs. 1 and 2), while all mock-infected cells developed large numbers of SGs in response to NaAs (Fig. 1).

**Fig. 3.**
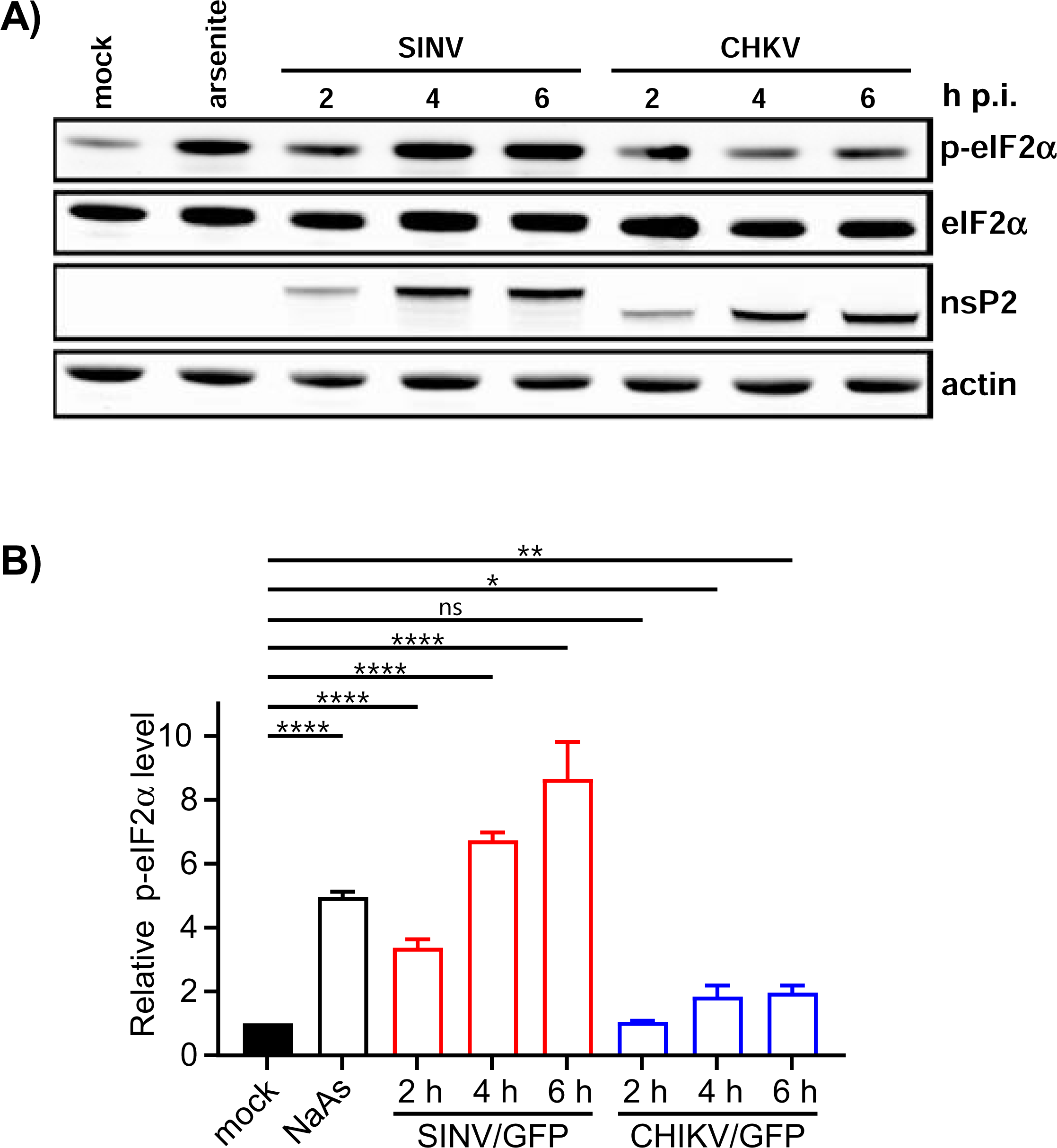
Phosphorylation of eIF2α is induced more efficiently by replication of SINV than CHIKV. A) NIH 3T3 cells in 6-well Costar plates (5×10^5^ cells/well) were infected with SINV/GFP and CHIKV/GFP at an MOI of 20 PFU/cell. At the indicated times p.i., cells were harvested. Mock-infected cells were treated with 0.75 mM NaAs for 45 min. Cell lysates were analyzed by WB using indicated antibodies and corresponding secondary antibodies, labeled with the infrared dyes. Membranes were scanned on the Odyssey Imaging System. B) Band intensities were determined in Empiria Studio 2.2. The p-eIF2α band intensities were first normalized to the total eIF2α and then to the normalized level of p-eIF2α in mock-infected cells. Means and SD are indicated. The significance of differences was determined by one-way ANOVA with the two-stage step-up method of Benjiamini, Krieger and Yekutieli test (****P ≤ 0.0001, **P ≤ 0.01, *P ≤ 0.05, ns ≥ 0.05; n = 3).

The data presented demonstrate that SINV and CHIKV infections alone are not efficient in inducing SG formation. Moreover, the replication of these viruses renders cells incapable of responding with SG assembly when treated with the well-characterized inducer NaAs. Given that SINV is a stronger activator of eIF2α phosphorylation, it likely possesses a more robust mechanism to suppress SG assembly.

### Replication of G3BP-independent SINV mutants

Our previous study and those of other research groups demonstrated that cellular G3BP1/2 proteins directly interact with short repeating peptides located in the C-termini of SINV-, SFV-, and CHIKV-specific nsP3 HVDs (28, 30, 31, 33). G3BPs play essential roles in promoting viral replication, and when the expression of both G3BPs is knocked out (*G3bp* dKO cells), this strongly affects SINV and SFV replication and completely abolishes replication of CHIKV (20, 33). Since SINV remains viable in NIH 3T3 *G3bp* dKO cells, we utilized this virus as an experimental system to further investigate the role of nsP3 HVD-G3BP interactions in inhibiting SG assembly.

In our initial experiments, we designed two SINV variants with mutated nsP3 HVDs. One variant, SINV/nsP3mut-GFP, had F-to-E mutations in both G3BP-binding repeat elements and another upstream peptide that shares some similarity with the canonical G3BP-binding motif (Fig. 4A). This upstream peptide could potentially facilitate additional weak interactions between the HVD and G3BPs. The other variant, SINV/nsP3del-GFP, had the entire repeat-encoding fragment, including a potential third G3BP-binding motif, deleted (Fig. 4A). Previous studies have shown that inserting GFP into the HVD of SINV nsP3 has only a minor negative effect on viral replication rates (35). Therefore, the parental construct and both mutants contained this GFP-coding sequence in their HVDs. The expression of nsP3-GFP fusion was used to analyze the formation of nsP3 complexes and their compartmentalization in virus-infected cells.

**Fig. 4.**
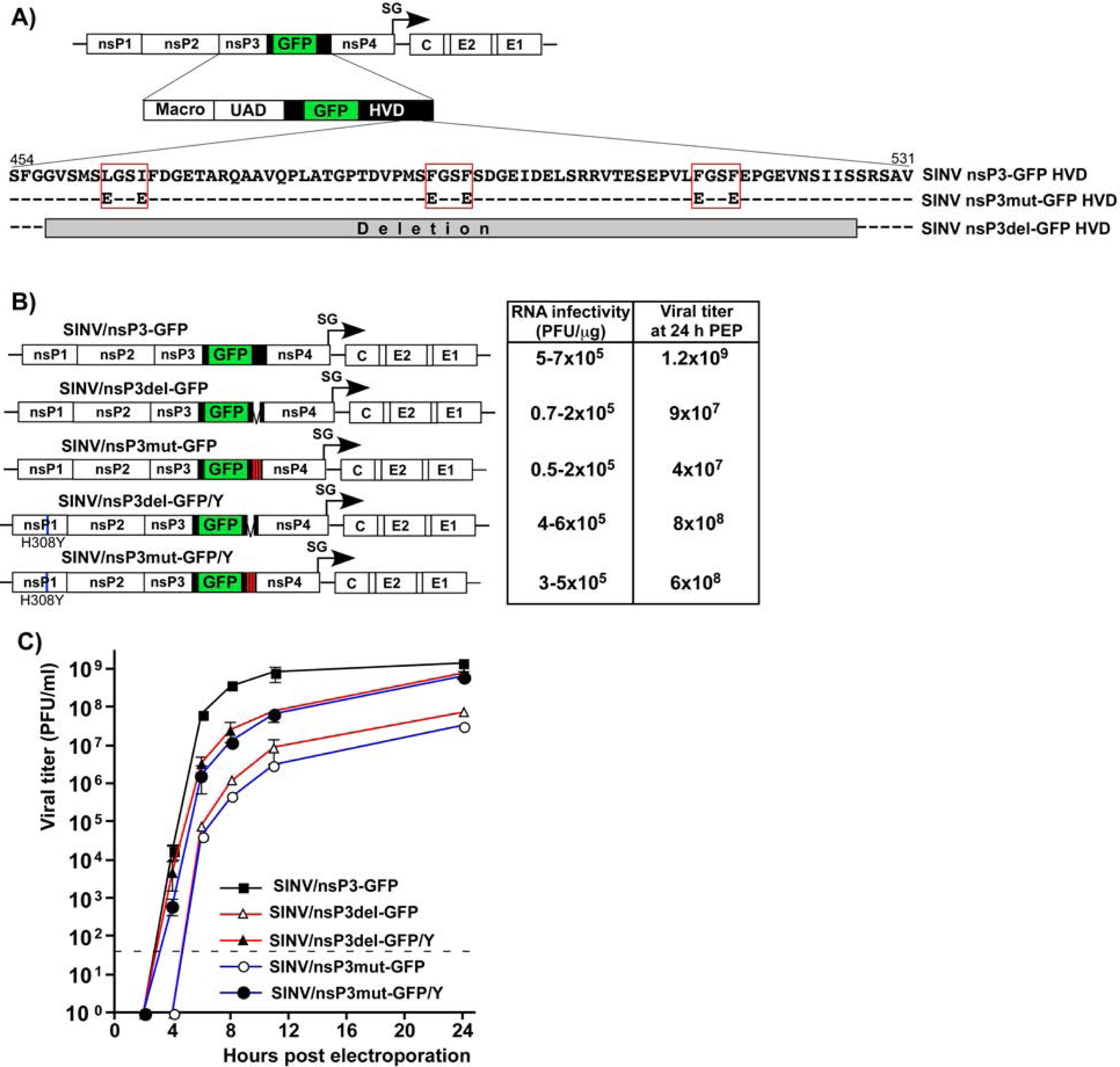
SINV/nsP3-GFP variants with mutated or deleted G3BP-binding sites in nsP3 HVD are viable but replicate less efficiently than the parental virus. A) The schematic presentation of the coding strategy of SINV/nsP3-GFP G RNA, the domain structure of nsP3-GFP, and mutations and deletions introduced into HVD. Red boxes indicate positions of the G3BP-binding sites. B) The schematic presentation of the G RNAs of SINV variants with mutated HVDs. BHK-21 cells were electroporated by 3 µg of the *in vitro*-synthesized G RNAs of these variants to generate viral stocks and analyze RNA infectivities in ICA. C) BHK-21 cells were electroporated by 3 µg of the *in vitro*-synthesized G RNAs. At the indicated time post electroporation, media were replaced, and viral titers were assessed by plaque assay on BHK-21 cells.

In the infectious center assay (ICA), the *in vitro*-synthesized RNAs of the designed SINV/nsP3mut-GFP and SINV/nsP3del-GFP showed lower infectivity compared to the parental SINV/nsP3-GFP (Fig. 4B). However, the differences were less than 10-fold, indicating that the designed mutants were viable and not second-site revertants. Viral titers at various time points post electroporation of BHK-21 cells were also 2-3 orders of magnitude lower than those of the parental SINV/nsP3-GFP. These observed differences in ICA and viral titers were expected based on our previous finding that HVD-G3BP complexes function in vRC assembly and recruitment of G RNA into replication (20).

The electroporation-derived stocks of the mutants, but not the parental SINV/nsP3-GFP, displayed heterogeneity in plaque size, and a small percentage of larger plaques indicated viral evolution towards more efficiently replicating phenotype. Therefore, the pools of rescued mutants were enriched with evolved, better replicating variants through three additional passages on BHK-21 cells. Large plaques were randomly selected, and the nsP-coding fragments of the genomes of plaque-purified mutants were sequenced. No additional changes were found in nsP2, nsP4, or the mutated nsP3, but a reproducible H308Y substitution was detected in SINV nsP1. This mutation was then introduced into the cDNAs of the originally designed SINV/nsP3mut-GFP and SINV/nsP3del-GFP. The presence of the H308Y mutation enhanced the infectivity of the *in vitro*-synthesized RNAs and the replication rates of the mutant viruses (Figs. 4B and C). The nsP1-specific mutation stabilized the designed SINV/nsP3del-GFP/Y and SINV/nsP3mut-GFP/Y variants, and no further evolution was detected in subsequent experiments. Since the adaptive H308Y mutation was not in the nsP3 HVD, it is highly unlikely that it affected HVD interactions with G3BPs and SG formation. Therefore, the designed variants with the H308Y mutation were used in the following experiments, and the compensatory effect of the nsP1-specific mutation was not further investigated. Thus, despite lacking G3BP-binding sites in their HVDs, the SINV variants were viable and became stable after acquiring a single mutation in nsP1.

### Mutated HVDs do not form complexes with G3BPs

Next, we confirmed that the mutated HVDs had lost the ability to bind G3BPs. The mutated and parental SINV nsP3 HVDs were fused with Flag-GFP and cloned into VEEV replicons under the control of the SG promoter (Fig. 5A). These replicons were then packaged into infectious viral particles, and NIH 3T3 cells were infected at the same MOI. At 3 h p.i., HVD-bound protein complexes were isolated using anti-Flag MAb magnetic beads and analyzed by WB using G3BP1-specific antibodies. The co-immunoprecipitation samples of both mutated HVD fusions showed no presence of G3BP (Fig. 5B), leading us to conclude that the mutated HVDs were no longer capable of binding murine G3BPs.

**Fig. 5.**
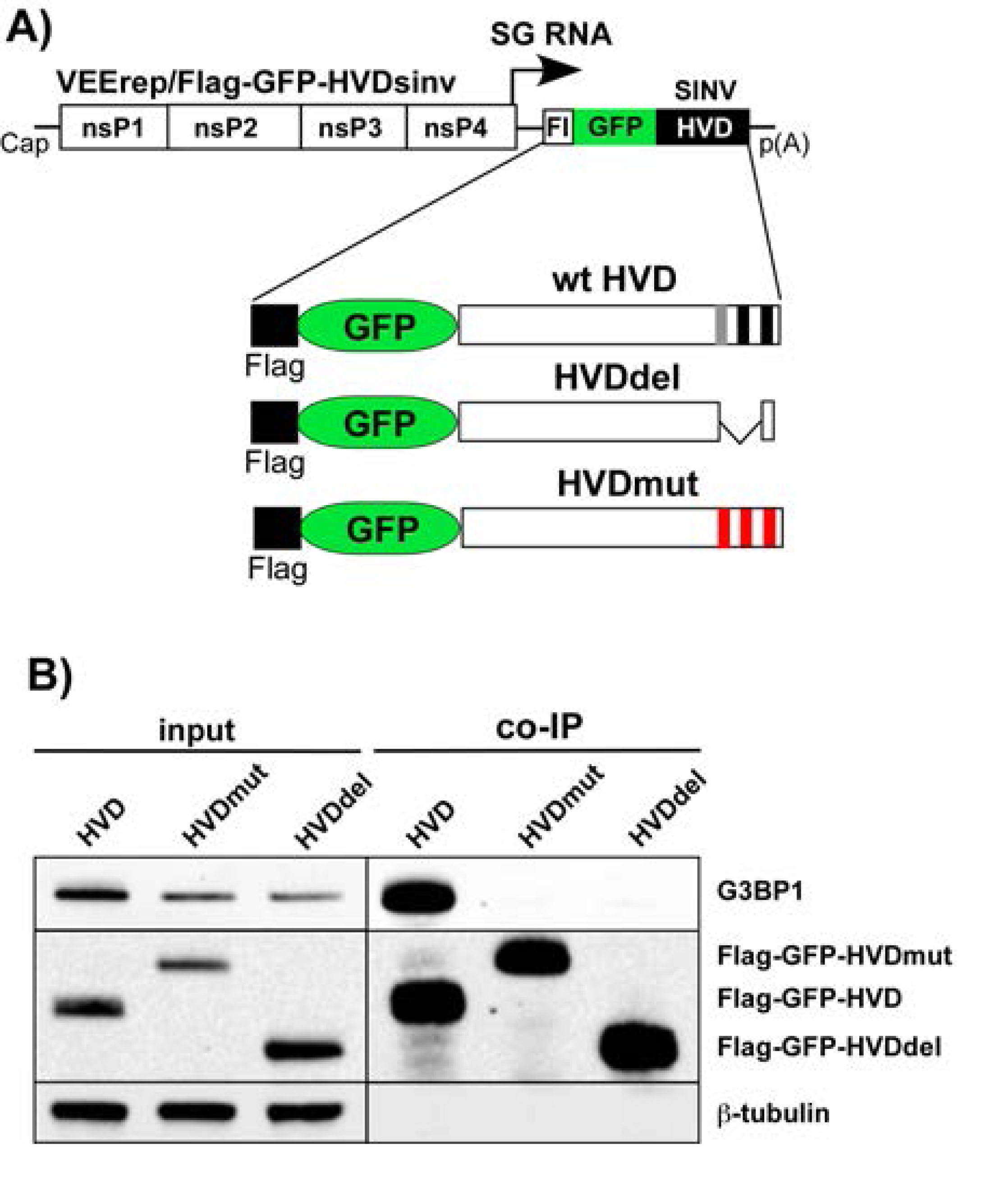
Mutated SINV nsP3 HVDs do not bind G3BPs. A) The schematic presentation of VEEV replicon and encoded Flag-GFP-HVDsinv fusions. B) NIH 3T3 cells were infected with VEEV replicons encoding indicated fusions at the same MOI. At 3 h p.i., cells were lysed, protein complexes were isolated using anti-Flag magnetic beads and analyzed by WB using Flag– and G3BP1-specific antibodies and infrared dye-labeled secondary antibodies. Images were acquired on Odyssey Imaging System.

Next, we investigated the effects of modifications in the G3BP-binding fragment of SINV HVD on the distribution and composition of nsP3 complexes formed during viral replication. Cells were infected with SINV/nsP3-GFP, SINV/nsP3del-GFP/Y, and SINV/nsP3mut-GFP/Y, and the distribution of nsP3-GFP and G3BPs was assessed. Unlike the parental SINV/nsP3-GFP, the replication of both viruses with mutated HVDs had no effect on the diffuse distribution of G3BPs, which remained similar to that found in uninfected cells (Fig. 6 for G3BP1 and data not shown for G3BP2). These findings demonstrated that the introduced modifications rendered SINV nsP3 HVDs incapable of interacting with G3BPs and forming nsP3-G3BP complexes during viral replication. Importantly, staining of cells with antibodies against the SG marker TIAR showed the absence of SGs at 6 h p.i. with SINV variants encoding mutated HVDs (Fig.6).

**Fig. 6.**
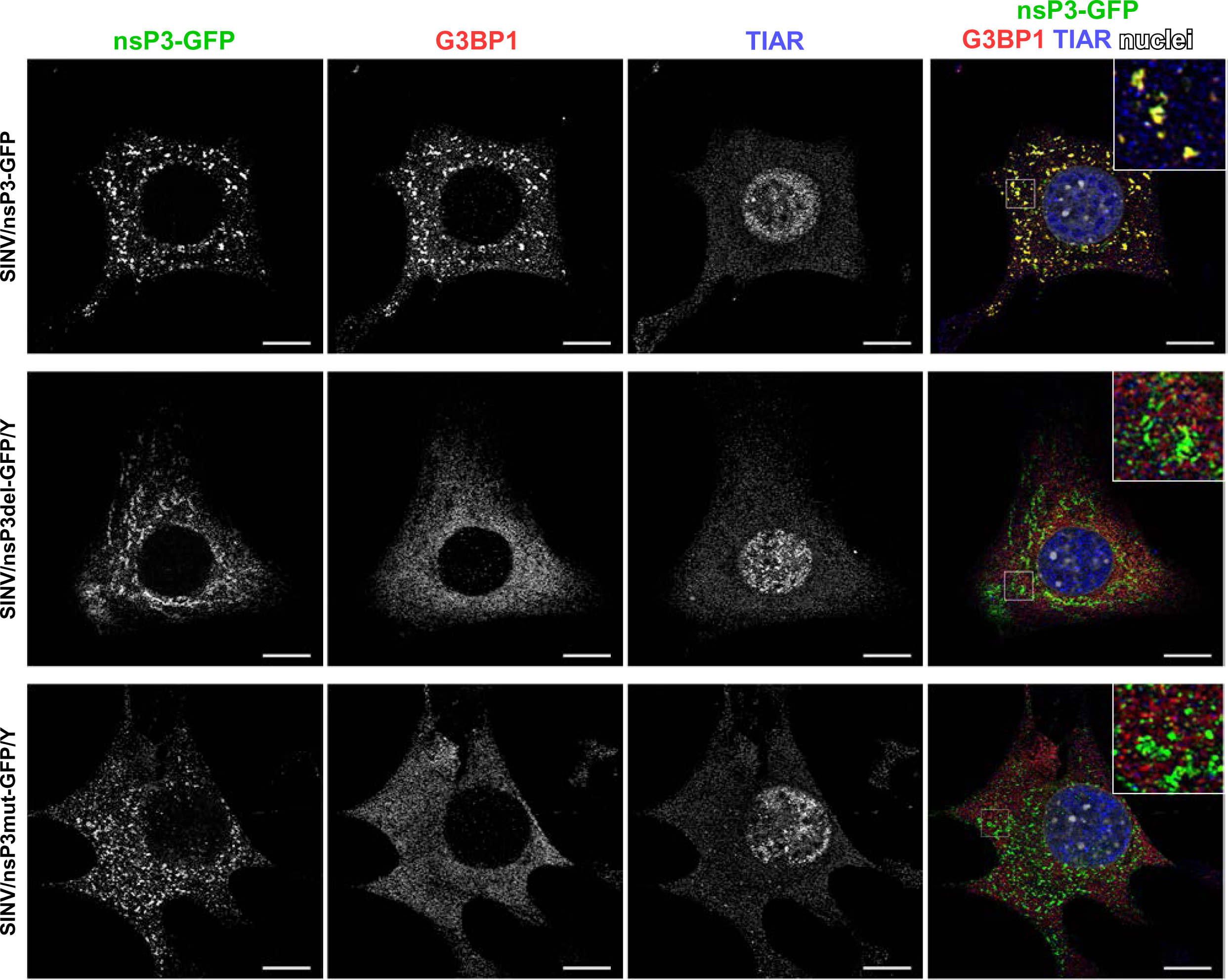
The designed viral variants with mutated HVD do not accumulate G3BP in nsP3 complexes and do not induce SG formation in response to their replication. NIH 3T3 cells were infected with the indicated viruses at an MOI of 20 PFU/cell. At 6 h p.i., they were fixed and immunostained with the antibodies specific to G3BP1 and TIAR. Bars correspond to 10 μM.

### The inability of cells to express G3BPs and form SGs did not have any positive effects on the replication of SINV variants with mutated nsP3 HVDs

We then used HVD mutants to investigate viral interference with SG formation and its role in SINV replication. They lacked G3BP-binding motifs in their HVDs, and their replication had no effect on G3BP distribution (Fig. 6). According to the prevailing hypothesis, such mutants could not sequester G3BPs into nsP3 complexes and interfere with SG development. Therefore, it was reasonable to expect that they would replicate less efficiently in the parental, SG-competent NIH 3T3 cells compared to their *G3bp* dKO derivative, which is unable to form SGs even in response to NaAs treatment (Fig. 7).

**Fig. 7.**
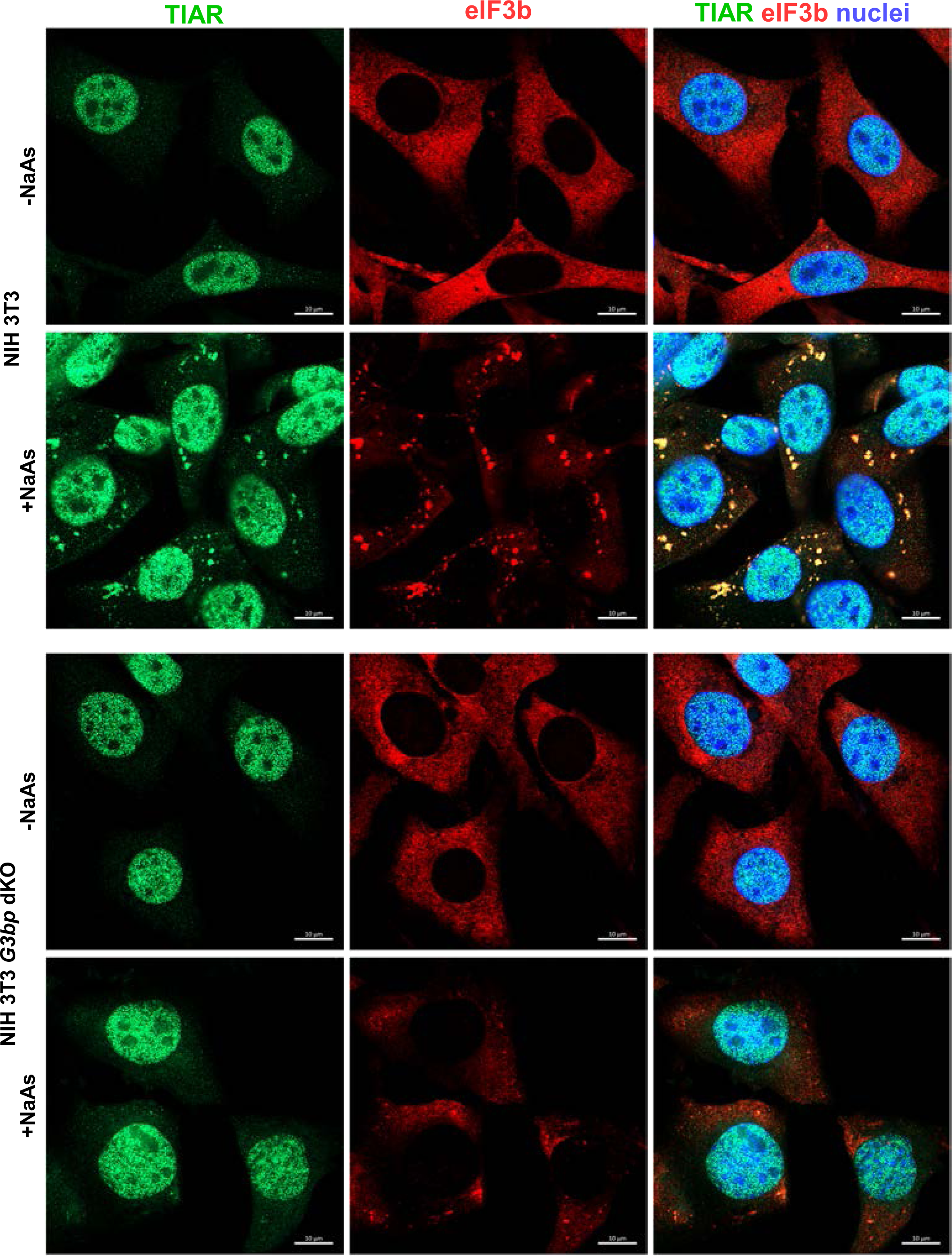
NIH 3T3 *G3bp* dKO cells do not respond to NaAs treatment by SG formation. *G3bp* dKO and parental NIH 3T3 cells were treated with 0.75 mM NaAs for 45 min or remained mock-treated. Then they were fixed and immunostained with the antibodies specific to SG components Bars correspond to 10 μM.

For these experiments, the viruses were modified. To eliminate any potential effects of GFP insertion on nsP3 function, all the newly designed constructs lacked GFP in their nsP3 HVDs (Fig. 8A). However, the GFP gene was inserted into the genomes of SINV/nsP3del/GFP/Y, SINV/nsP3mut/GFP/Y, and SINV/GFP/Y under the control of additional SG promoters. GFP expression facilitated monitoring the spread of the viral mutants, which replicate less efficiently than the control virus in cell culture. All the constructs also contained the previously identified H308Y mutation in nsP1. The viruses were generated by electroporation of *in vitro*-synthesized RNAs into BHK-21 cells. First, we compared their abilities to form plaques on NIH 3T3 and NIH 3T3 *G3bp* dKO cell lines (Fig. 8B). As expected, SINV/GFP/Y, expressing wild type nsP3 HVD, formed large plaques on NIH 3T3 cells.

**Fig. 8.**
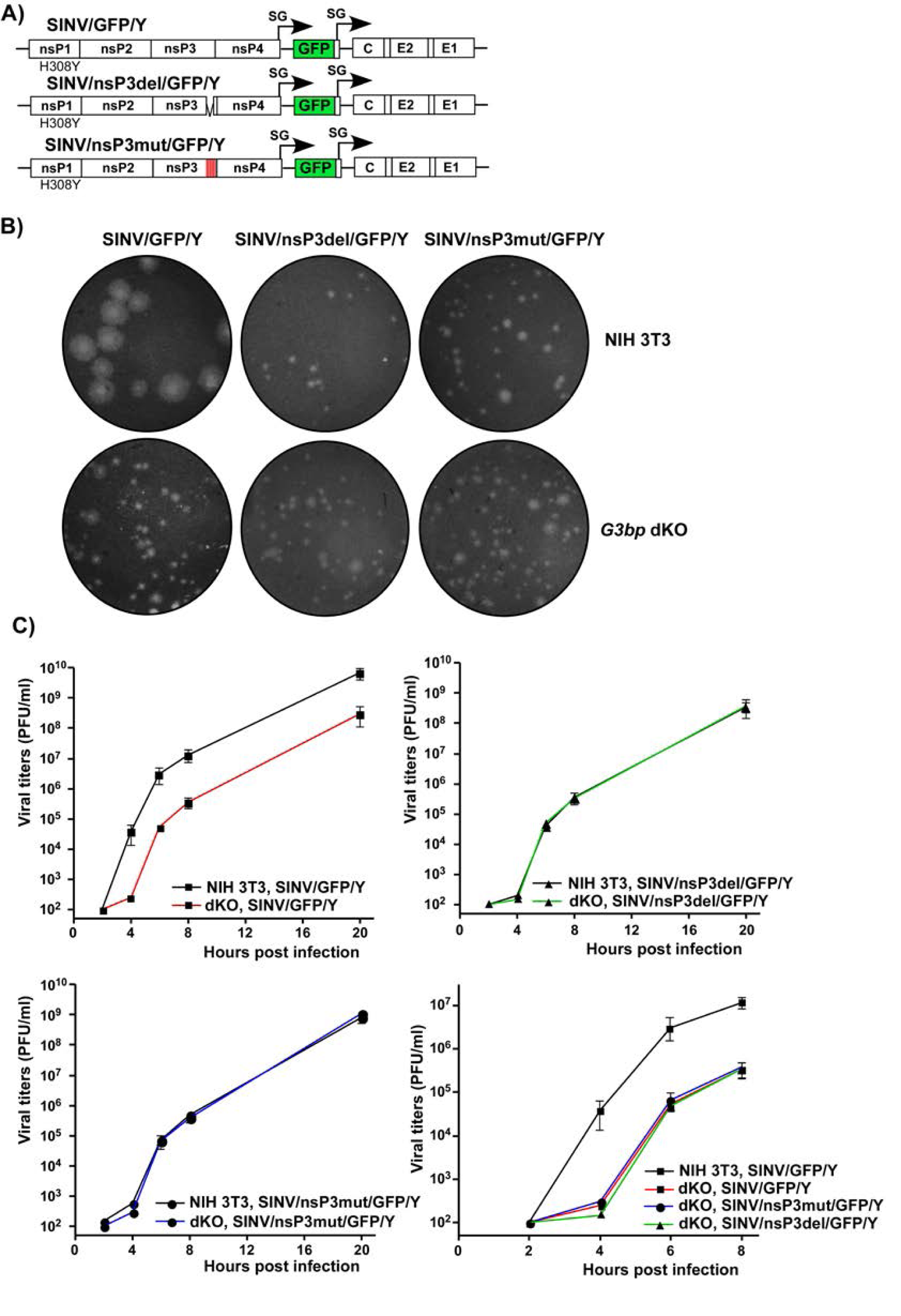
Lack of G3BP expression and inability of cells to form SGs does not stimulate replication of SINV variants with mutated HVDs. A) The schematic presentation of G RNAs of SINV variants with mutated HVDs. B) Equal numbers of *G3bp* dKO and parental NIH 3T3 cells were seeded into 6-well Costar plates and used in the plaque assay done on the indicated variants. Cells were fixed by paraformaldehyde at 2 days p.i. and plaques were immunostained by crystal violet. C) *G3bp* dKO and parental NIH 3T3 cells were seeded into 6-well Costar plates (5×10^5^ cells/well) and infected with the indicated variants at an MOI of 0.01 PFU/cell. At the indicated times p.i., media were replaced and viral titers were determined by plaque assay on BHK-21 cells. The experiment was repeated 3 times with the identical results.

However, its plaques on the *G3bp* dKO cell line were dramatically smaller due to the lack of G3BPs required for efficient vRC assembly and function. The sizes of plaques produced by SINV/nsP3del/GFP/Y and SINV/nsP3mut/GFP/Y mutants in both NIH 3T3 and *G3bp* dKO cells were consistently small and indistinguishable from those of SINV/GFP/Y produced on *G3bp* dKO cells (Fig. 8B). The foci of GFP-positive cells around plaques formed by the mutants were also small (data not shown). This was an indication that the small plaque size was primarily determined by inefficient viral replication and spread rather than a lower ability of the mutants to induce cell death and cause cytopathic effect (CPE). In conclusion, these experiments demonstrated that the absence of structural SG components (G3BPs) in dKO cells did not stimulate viral spread and CPE development.

Next, SINV/nsP3del/GFP/Y, SINV/nsP3mut/GFP/Y, and SINV/GFP/Y variants were compared in terms of replication rates in *G3bp* dKO and parental NIH 3T3 cells (Fig. 8C). Consistent with the plaque size data, SINV/GFP/Y replicated more efficiently in NIH 3T3 than in dKO cells. At any time p.i., its titers in NIH 3T3 cells were 50-100-fold higher than those in the dKO counterpart. In contrast, both mutants replicated with equal efficiency in NIH 3T3 and *G3bp* dKO cells. Moreover, their replication rates were identical to those of SINV/GFP/Y in the dKO cell line. These data clearly demonstrated that the inability of cells to express G3BPs and form SGs had no stimulatory effect on the replication of SINV mutants encoding nsP3 HVDs incapable of interacting with G3BPs.

### SINV mutants inhibit SG assembly even during NaAs treatment

In the experiments presented in Fig. 6, we observed that despite the mutant nsP3s did not interact with G3BPs, infections with mutant viruses did not lead to SG development. However, this does not prove that the mutant viruses are incapable of preventing SG formation in response to chemical stress inducers, such as NaAs. To investigate this further, NIH 3T3 cells were infected with SINV/nsP3del/GFP/Y, SINV/nsP3mut/GFP/Y, and SINV/GFP/Y and then treated with NaAs at 6 h p.i. Exposure of mock-infected cells to NaAs induced the formation of large G3BP-containing SGs in all cells. However, less than 1% of the cells infected with SINV encoding either wild-type nsP3 or nsP3 with HVDs lacking the G3BP-binding motifs exhibited the presence of a few SGs (Fig. 9). The same result was obtained on BHK-21 cells (data not shown). Thus, despite the inability to sequester G3BP into nsP3 complexes, replication of SINV HVD mutants prevented SG formation in response to NaAs treatment.

**Fig. 9.**
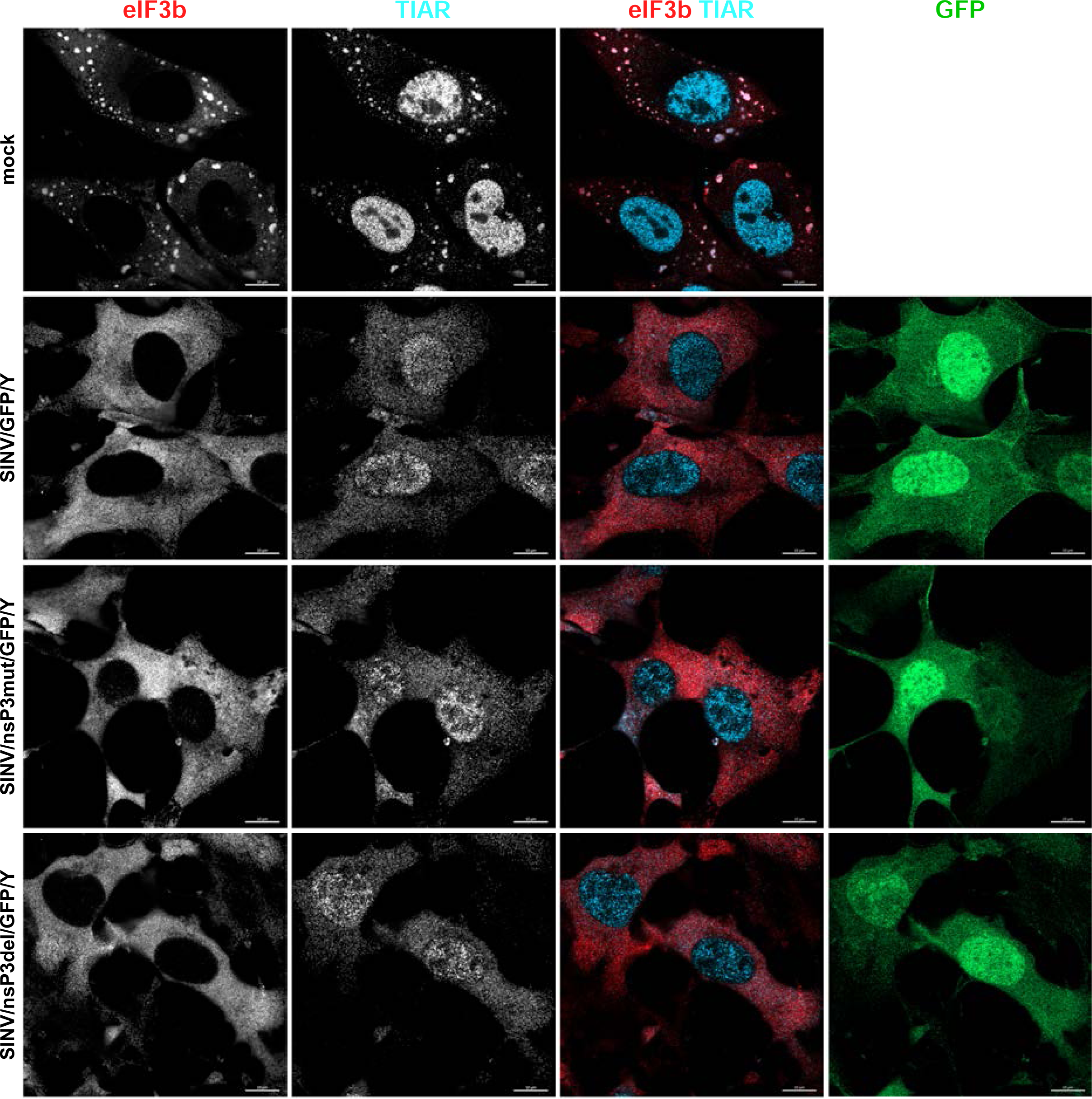
Cells infected with either SINV HVD mutants or parental virus are incapable of forming SGs in response to NaAs treatment. NIH 3T3 cells were infected with the indicated viruses at an MOI of 10 PFU/cell. In 6 h, infected and mock-infected cells were exposed to 0.75 mM NaAs for 45 min, fixed, and immunostained with antibodies specific to the indicated components of SGs. Bars correspond to 10 μM.

### Stable cell lines expressing high levels of either nsP3 or its HVD form SGs upon NaAs treatment

Previous experiments demonstrating the inhibitory functions of nsP3 in SG assembly relied on transient overexpression of nsP3 alone (29, 32). However, we found that the transfection reagents strongly and nonspecifically inhibit SG formation (data not shown), making the results difficult to interpret. Therefore, we established stable cell lines expressing either full-length SINV nsP3 or its HVD fused with GFP. Clones with high levels of nsP3 fusion produced large nsP3-GFP aggregates and were excluded from further analysis, and only those with lower levels of expression and diffuse distribution of GFP fusion proteins were used. It is worth noting that the selected clones expressed the GFP-nsP3 fusion at higher levels than that observed in NIH 3T3 cells at 6 h p.i. with SINV/nsP3-GFP (Fig. 10A). At this time p.i., infected cells become incapable of SG development even in response to NaAs treatment (Fig. 1). However, despite the higher concentration of GFP-nsP3 in the stable cell lines, the TIAR– and eIF3b-positive SGs were formed in response to NaAs as efficiently as in naïve NIH 3T3 cells (Fig. 10B). This strongly indicates that neither the HVD nor other domains play critical roles in making cells resistant to SG formation. Additionally, similar results were obtained with NIH 3T3 cells stably expressing GFP fused with full-length CHIKV nsP3: they also formed SGs in response to NaAs (data not shown). Thus, GFP-nsP3 expression in stable cell lines did not interfere with SG formation. In the cell lines expressing GFP-HVD, this fusion protein noticeably interfered with SG generation during NaAs treatment (Fig. 10B). However, the latter fusion was also expressed at very high concentrations and its negative effect was dependent on the levels of expression.

**Fig. 10.**
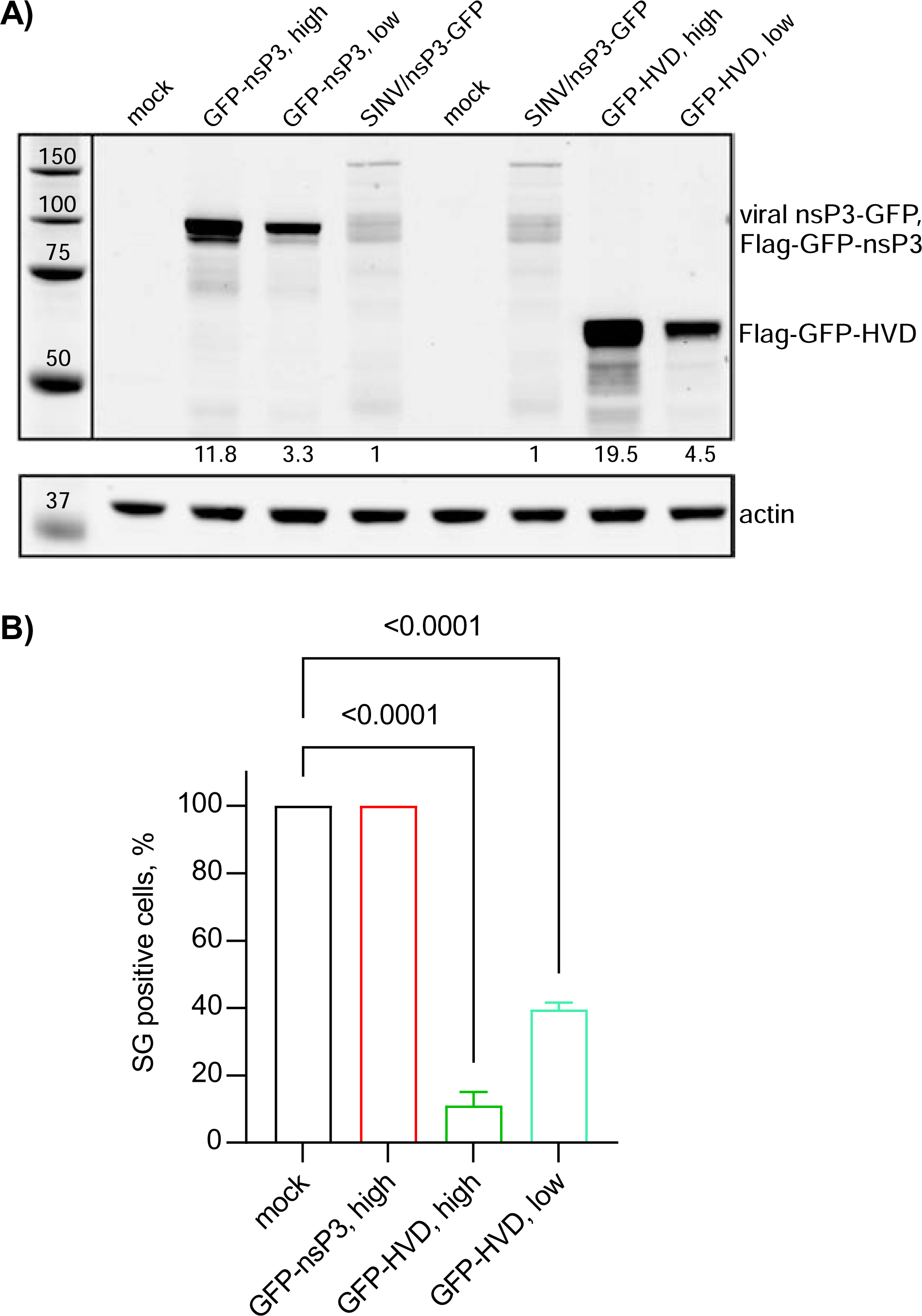
Stable cell lines expressing SINV nsP3 and HVD fusions remain capable of forming SGs in response to NaAs treatment. A) Stable cell lines of NIH 3T3 cells expressing Flag-GFP-nsP3 and Flag-GFP-HVD fusions were generated as described in Materials and Methods. Individual clones of GFP positive cells were analyzed by WB in terms of the expression of fusion proteins using GFP-specific antibodies. The levels of expression were compared to that of nsP3-GFP in NIH 3T3 cells at 6 h p.i. with SINV/nsP3-GFP. B) The indicated clones of stably expressing cells were treated with 0.75 mM NaAs for 45 min, fixed and immunostained with antibodies specific to SG components (eIF3b and TIAR). A percentage of GFP-positive cells containing SGs was determined (∼100 cells per experiment). Means and SD are indicated. The significance of differences was determined by one-way ANOVA with the Dunnett test (n = 3).

From these experiments, two conclusions were drawn. Firstly, neither SINV nsP3 HVD nor the entire SINV and CHIKV nsP3s could solely act as primary inhibitors of SG development. Thus, during the expression at low concentrations achieved by virus replication at early times p.i., nsP3’s interaction with G3BPs may probably have a supporting role in SG inhibition, but this is not the major factor. Secondly, the ADP-ribosylhydrolase function of the nsP3-specific macro domain of SINV and CHIKV nsP3s is also insufficient to inhibit SG formation, and NIH 3T3 cells stably expressing SINV and CHIKV nsP3s are not resistant to SG formation when exposed to NaAs. To rule out the possibility that GFP fusion to the N-terminus of nsP3 could interfere with the macro domain’s ADP-ribosylhydrolase activity, GFP was also fused to the C-terminus of the protein. The NIH 3T3 cells stably expressing this fusion remained fully capable of developing SGs when exposed to NaAs (data not shown).

### Transcriptional and translational shutoffs interfere with SG assembly in alphavirus-infected cells

One of the common characteristics of the OW alphaviruses is a rapid induction of transcriptional and translational shutoffs (36). They fully develop within 4-6 h p.i. and coincide with the acquisition of cell resistance to SG formation. To investigate the possible role of transcription inhibition in developing resistance of the infected cells to SG formation, we mimicked alphavirus-induced transcriptional shutoff by treating NIH 3T3 cells with Actinomycin D (Act D) (Fig. 11). NIH 3T3 cells were incubated in Act D-supplemented media for 1, 2, and 4 h before being exposed to NaAs. Immunostaining for SG markers clearly demonstrated that the cells already poorly formed SGs in response to NaAs after 2 h of Act D exposure. After 4 h of Act D treatment, NaAs was no longer able to induce SG assembly at all. This suggested that alphavirus-induced transcription inhibition may be critically involved in interference with SG development even in response to the chemical inducer.

**Fig. 11.**
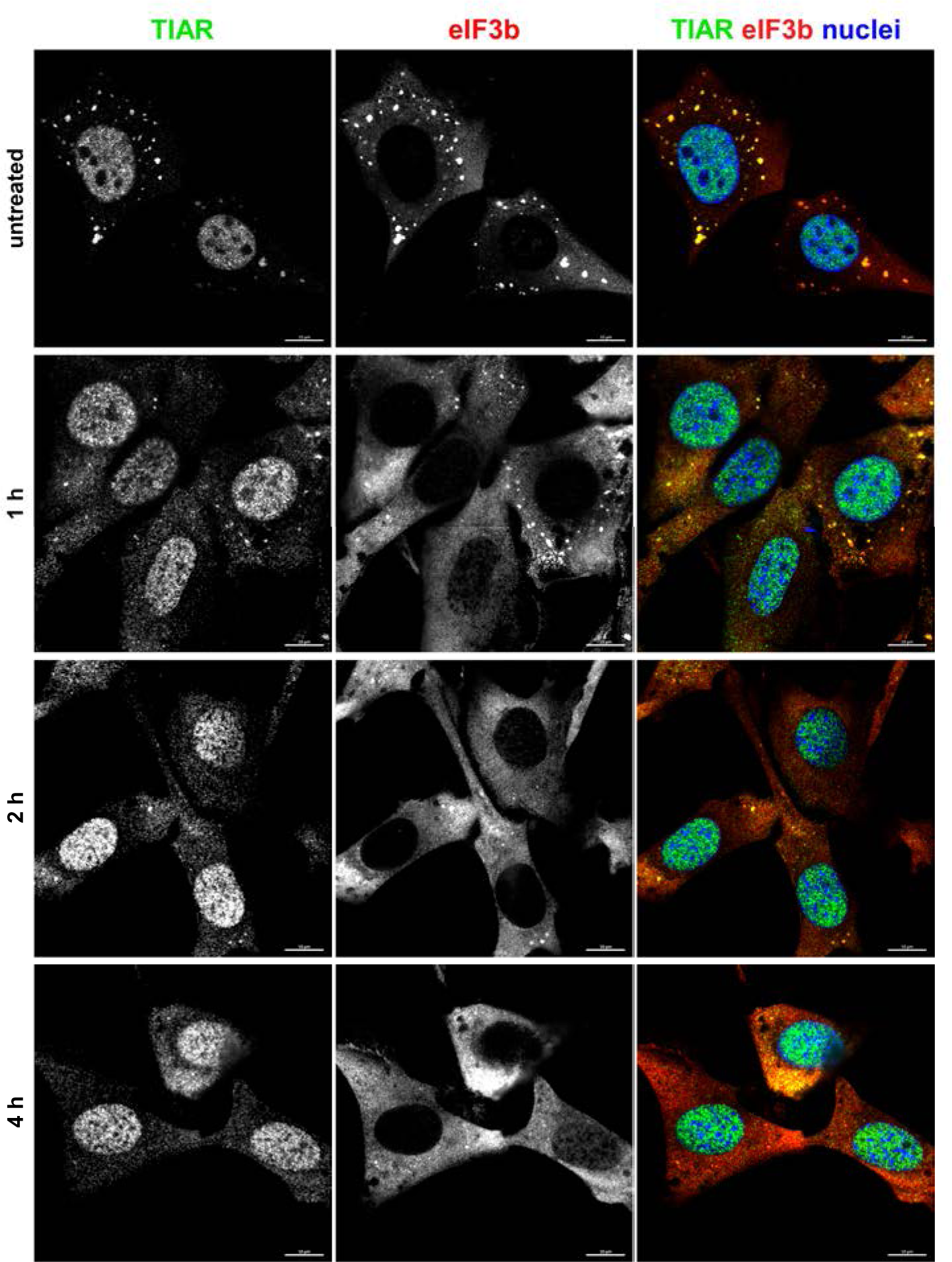
Transcriptional shutoff induced by ActD rapidly blocks formation of SGs in response to NaAs treatment. NIH 3T3 cells were treated with ActD for indicated times and then exposed to 0.75 mM NaAs for 45 min. Ultimately, cells were fixed and immunostained with antibodies specific to the indicated components of SGs. Bars correspond to 10 μM.

In previous studies, we generated several SINV mutants that exhibited reduced cell inhibitory functions. One of them, SINV with the P726G mutation in nsP2 (SINV/G/GFP), was less efficient in inhibiting both cellular transcription and translation (36, 37). Consequently, we infected NIH 3T3 cells with wt SINV/GFP and SINV/G/GFP carrying the mutated nsP2. No SGs were observed in cells infected with SINV/GFP at 6 h p.i., and treatment of the infected cells with NaAs also failed to induce SG development (data not shown and Fig. 12B). However, over 30% of cells infected with SINV/G/GFP formed SGs after NaAs treatment (Figs. 12A and B).

**Fig. 12.**
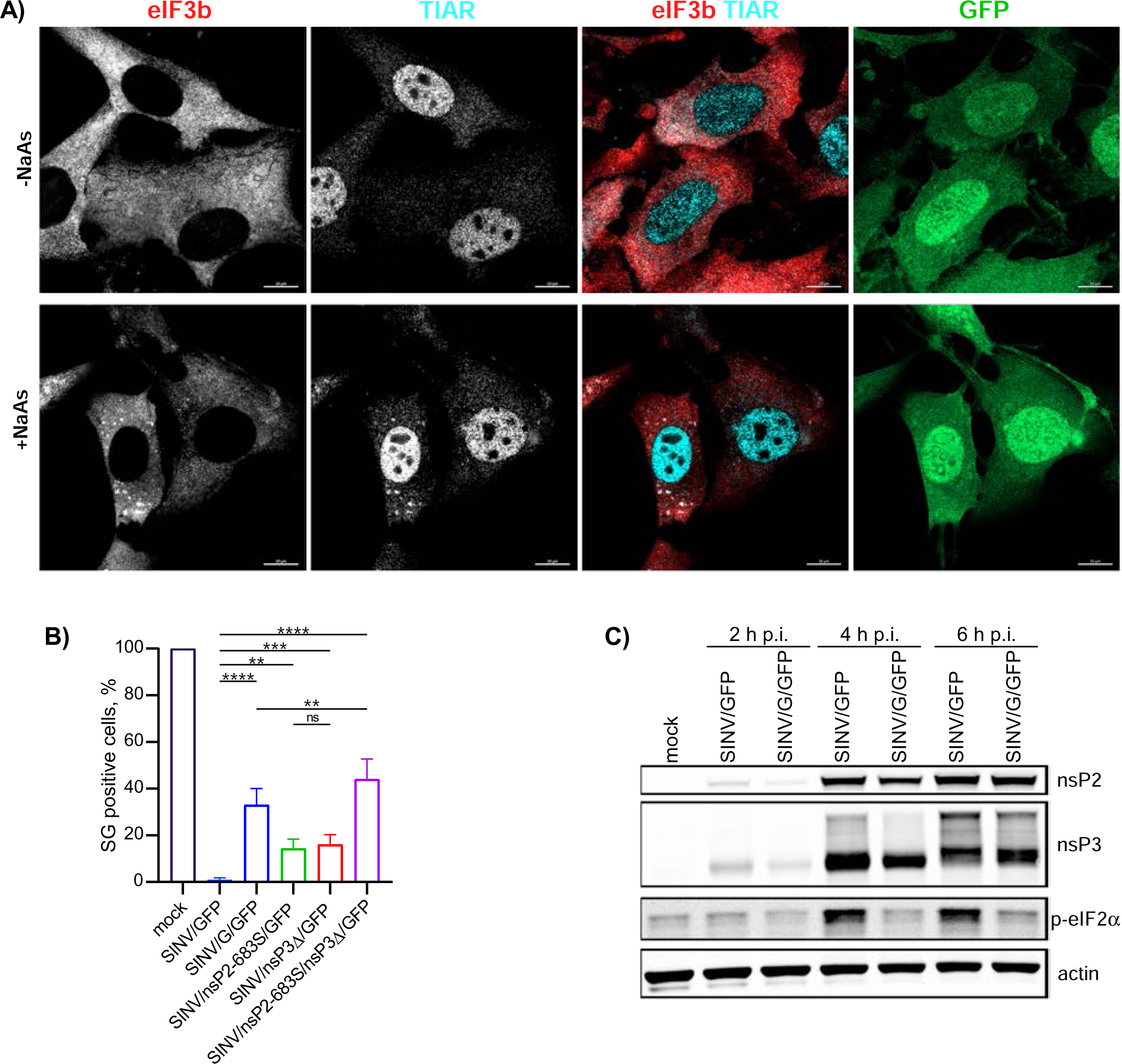
The inabilities of viral mutants to develop transcriptional or translational shutoffs make them incapable of blocking SG development in response to NaAs treatment despite the accumulation of wt nsP3. A) NIH 3T3 cells were infected with SINV/G/GFP at an MOI of 10 PFU/cell. At 6 h p.i., they were treated with 0.75 mM NaAs for 45 min or remained mock-treated. Cells were fixed, immunostained with antibodies specific to the indicated SG markers and analyzed by confocal microscopy. Bars correspond to 10 μM. B) NIH 3T3 cells were infected with the indicated viral variants (see the text for details) at an MOI of 10 PFU/cell, fixed and immunostained with TIAR– and eIF3b-specific antibodies to detect SGs. A percentage of GFP-positive cells containing SGs was determined (∼100 cells per experiment). Means and SD are indicated. The significance of differences was determined by one-way ANOVA with the two-stage step-up method of Benjiamini, Krieger and Yekutieli test (****P ≤ 0.0001, ***P≤0.001, **P ≤ 0.01, ns ≥ 0.05; n = 3). C) NIH 3T3 cells in 6-well Costar plates were infected with SINV/G/GFP and parental SINV/GFP at an MOI of 10 PFU/cell, and harvested at the indicated times p.i. Cell lysates were analyzed by WB for accumulation of viral nsPs and p-eIF2α.

The inability of SINV/G/GFP to block SG formation correlated with its reduced ability to induce phosphorylation of PKR and eIF2α [(21) and Fig. 12C] that is required for SG development. Notably, this mutant virus produced both nsP2 and, more importantly, wt nsP3 at levels comparable to those found in the cells infected with parental SINV/GFP (Fig. 12C).

In addition to their reduced ability to inhibit cellular transcription and translation, the SINV/G variant also exhibits decreased rates of RNA and virus replication (15), and this could be an alternative explanation for its inability to interfere with SG induction. Consequently, we investigated three additional mutants with replication rates similar to those of the parental wt virus (36). SINV/nsP2-683S/GFP carried a mutation in nsP2 and was shown not to inhibit cellular transcription. SINV/nsP3Δ/GFP had a 6-aa-long deletion in nsP3 (Δ24-29) and inhibited cellular translation less efficiently than the wt virus. SINV/nsP2-683S/nsP3Δ/GFP, in turn, carried both mutations and did not inhibit both cellular transcription and translation. After infecting NIH 3T3 cells with these mutants for 6 h, we did not detect the appearance of SGs (data not shown). However, treatment of infected cells with NaAs resulted in SG generation (Fig. 12B). Approximately 15% of cells infected with either single mutant and around 40% of cells infected with the double mutant produced SGs in response to NaAs. Thus, we concluded that both virus-induced inhibition of transcription and translation are the critical determinants of the cells’ inability to assemble SGs in response to NaAs, and their effects appear to be additive.

To further investigate the hypothesis that inhibition of cellular transcription and translation affect SG development, we employed an additional experimental system. BHK-21 cells were transfected with a SINV replicon (SINrep/L/GFP/Pac) encoding GFP and puromycin acetyltransferase (Pac) under the control of different SG promoters. This replicon had a mutation (P726L) in nsP2, making it highly attenuated and allowing it to persistently replicate in cells lacking a functional type 1 IFN system (38). Importantly, SINrep/L/GFP/Pac encoded wt nsP3. BHK-21 cells were transfected with the *in vitro*-synthesized SINrep/L/GFP/Pac RNA and passaged in puromycin-containing media for 4 days. No SGs were detected in the replicon-carrying cells (Fig. 13A) and TIAR did not re-localize to the cytoplasm. Subsequently, the cells were treated with NaAs and immunostained for SG markers. Despite the presence of viral nonstructural proteins at the levels similar to those found in cells infected with SINV/GFP at 6 h p.i. (Fig. 13C), the replicon-containing Pur^R^ BHK-21 cells were able to form SGs in response to NaAs as efficiently as mock-infected cells (Fig. 13A). However, when the replicon-containing cells were superinfected with wt SINV/GFP, within 6 h p.i., they lost the ability to form SGs in response to NaAs treatment (Fig. 13A). The ability of superinfecting virus to replicate in the replicon-containing cells and block the SG development was confirmed by infecting them with SINV/nsP3-Cherry. nsP3-Chery complexes were readily detectable at 6 h p.i. (Fig. 13B). These findings provide additional evidence supporting the hypothesis that the inhibition of SG assembly in cells infected with SINV and potentially other Old World alphaviruses is primarily determined by the inhibition of cellular transcription and translation, rather than the expression of nsP3.

**Fig. 13.**
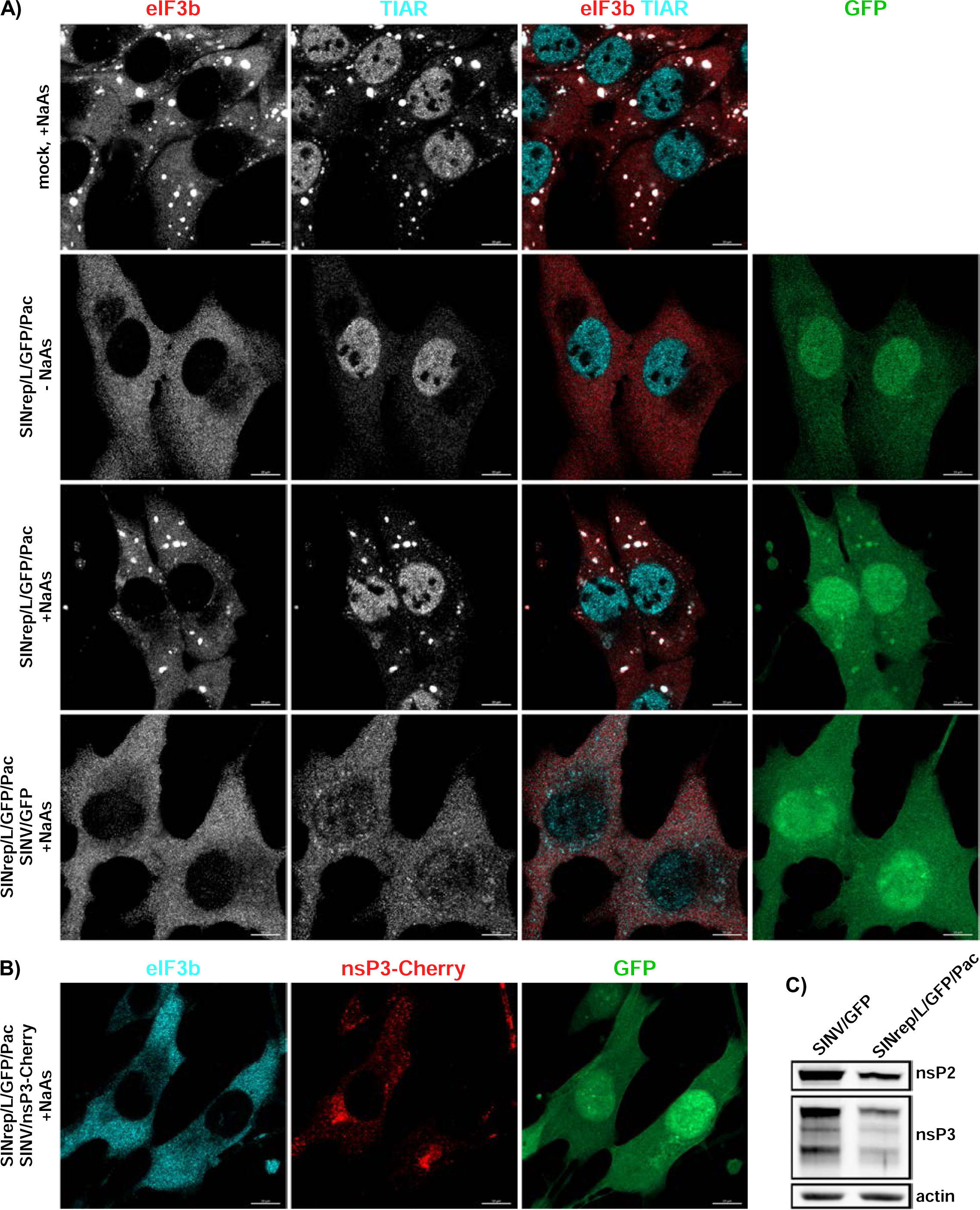
Persistent replication of SINV replicon does not inhibit SG formation during NaAs treatment. A) Naïve BHK-21cells, and cells carrying SINrep/L/GFP/Pac replicon were exposed to 0.75 mM NaAs for 45 min, fixed and immunostained with antibodies specific to SG components. The replicon-containing cells were also superinfected with SINV/GFP at an MOI of 10 PFU/cell and at 6 h p.i., treated with NaAs and immunostained with the same antibodies. Bars correspond to 10 μM. B) To show that SINrep/L/GFP/Pac-containing cells are readily superinfected with homologous virus, they were infected with SINV/nsP3-Cherry for 6 h, treated with NaAs as above, and distributions of nsP3-Cherry and the SG marker eIF3b were analyzed. C) WB analysis of nsP2 and nsP3 levels in the cells carrying SINrep/L/GFP/Pac and in those infected with SINV/GFP for 6 h. Membranes were scanned on the Odyssey Imaging System.

Previous studies have proposed that the efficient translation of viral SG RNA is involved in the inhibition of SG formation (28). The distinguishing characteristic of SINV replicons is that their SG RNAs are translated very inefficiently in the presence of p-eIF2α unless they contain a translational enhancer located at the beginning of the capsid-coding sequence (39). To test whether viral structural proteins and efficient translation of viral SG RNA play critical roles in SG formation, we infected BHK-21 cells with packaged SINVrep/GFP. The latter replicon expresses GFP 30-fold less efficiently than its counterpart containing the translational enhancer (21). However, similar to what we described earlier for SINV infection, less than 1% of cells were capable of forming SGs in response to NaAs treatment at 6 h p.i. with SINrep/GFP (Fig. 14). Therefore, it is unlikely that the efficient translation of SINV SG RNA significantly contributes to the inhibitory effect of SINV replication on SG assembly. However, we cannot completely rule out its minor contribution.

**Fig. 14.**
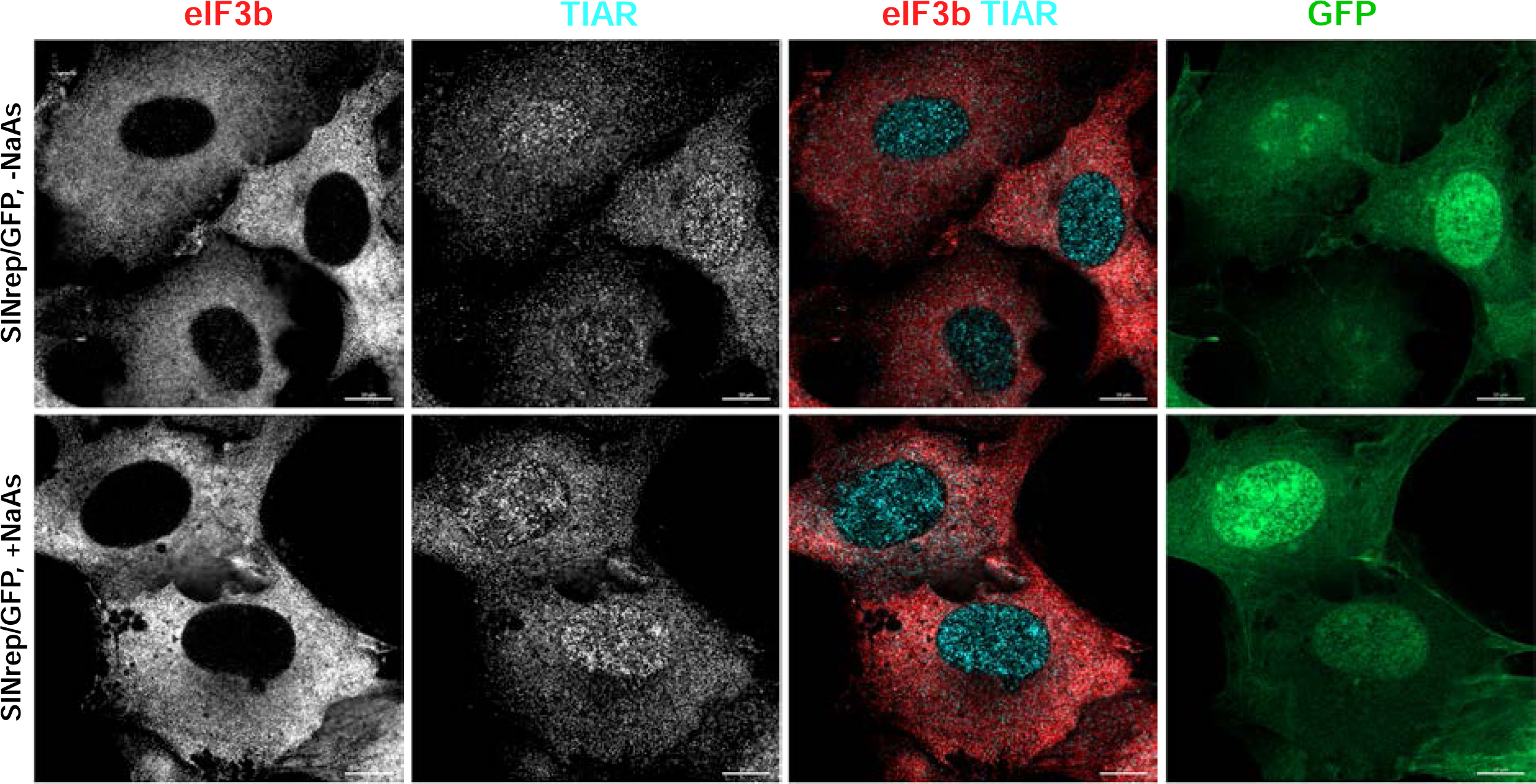
SINV replicons make cells resistant to SG formation. BHK-21 cells were infected with packaged SINrep/GFP replicon at an MOI of 10 inf.u/cell. At 6 h p.i., they were either treated with 0.75 mM NaAs for 45 min or mock-treated, fixed and immunostained with antibodies specific to SG markers. Bars correspond to 10 μM.

## Discussion

One of the key features of alphaviruses is their highly efficient replication in vertebrate cells. Following infection with the OW alphaviruses, such as SINV, SFV, and CHIKV, cells begin releasing viral progeny within 4-5 h p.i. By 16-24 h p.i., these cells typically demonstrate virus-induced CPE, characterized by profound morphological changes, loss of integrity, and cell detachment. During the first hours of viral replication, cells experience multiple changes in their biology, which may have both pro– and antiviral effects.

Replication of alphaviruses leads to the production of the dsRNA replication intermediates (18, 35). Additionally, vRCs can produce dsRNAs as nonspecific byproducts using cellular mRNA templates (40). Most dsRNAs are located inside membrane spherules (19, 40, 41). However, particularly during the early stages of infection, dsRNAs still can be sensed by cellular PRRs, such as RIG-I and MDA5 (22, 40). The presence of dsRNAs also activates PKR, and ultimately increases the levels of p-eIF-2α. This, in turn, leads to the accumulation of translationally stalled 48S initiation complexes, which are a prerequisite for the formation of SGs that accumulate translationally inactive mRNAs. However, during alphavirus infections, the development of SGs is rapidly blocked (28), and cells become resistant to SG induction even when exposed to potent chemical inducers such as NaAs. Consequently, both cellular and viral RNAs are not sequestered into SGs and may remain available for translation.

Previous studies have shown that the main components of SGs, G3BP1 and G3BP2, interact with nsP3 HVDs of the OW alphaviruses and eastern equine encephalitis virus (EEEV) (28, 30, 31, 33, 42). It was proposed that vRCs and large cytoplasmic nsP3 complexes sequester the entire pool of G3BP1/2, rendering these proteins unavailable for SG formation. These findings of nsP3-G3BP binding and the accumulation of G3BPs in nsP3 complexes provided a plausible explanation for the lack of SG formation during replication of at least some alphaviruses (28, 32). However, accumulating experimental data have suggested that nsP3 HVD-G3BP interactions are required for vRC formation and function rather than for the inhibition of SG development (20). In fact, the lack of nsP3-G3BP interaction in *G3bp* dKO cells renders CHIKV non-viable, and SINV replicates in this cell line several orders of magnitude less efficiently. Our new data also demonstrate that stable, high-level expression of the full-length SINV or CHIKV nsP3s, fused with GFP does not prevent SG assembly after exposure of the cells to NaAs. Expression of HVD alone can suppress SG development, but only at concentrations that are significantly higher than those of nsP3 achieved in virus-infected cells.

Importantly, SINV mutants containing mutations in nsP3 HVD that prevent interaction with G3BP still efficiently suppress SG assembly. Moreover, the absence of G3BP expression in *G3bp* dKO cells, which are incapable of building SGs, provides no detectable benefit for the replication of viral mutants lacking G3BP-binding sites in their nsP3 HVD. Thus, our data do not support a significant role for G3BP interactions with SINV and CHIKV nsP3 HVDs in the regulation of SG formation during viral replication.

Another recently proposed mechanism is based on the function of the macro domain of alphavirus nsP3 as an ADP-ribosylhydrolase (43–45). According to this hypothesis, the enzymatic activity of nsP3’s macro domain regulates the composition of SGs by reducing ADP-ribosylation of G3BP1 (29). However, as mentioned earlier, stable expression of the entire SINV or CHIKV nsP3-GFP fusions did not have noticeable effects on the ability of cells to form SGs after exposure to NaAs. Even though the expression levels of nsP3 fusions remained higher than those of nsP3 in virus-infected cells, these proteins could not interfere with SG formation. Therefore, it is difficult to expect that the ADP-ribosylhydrolase activity of the nsP3 macro domain plays a major role in downregulating SG assembly in infected cells. In this study, we also used a variety of SINV-based mutants SINV/G/GFP, and SINrep/L/GFP/Pac replicon, which expressed wt nsP3 at levels similar to that found in wt SINV-infected cells. Despite the presence of SINV nsP3 at levels relevant to wt virus infection, cells readily responded to NaAs treatment by forming SGs. This provides further evidence that nsP3-specific functions do not play major roles in the alphavirus-induced inhibition of SG assembly.

Our new data suggest that the inhibition of SG formation during replication of SINV and likely other alphaviruses is a complex process involving multiple components. Among them, the binding of G3BPs to CHIKV and SINV nsP3 HVDs, as well as the ADP-ribosylhydrolase activity of the nsP3 macro domain, do not significantly contribute to the inhibition of SG development, at least when nsP3 is expressed at biologically relevant levels. Instead, our data suggest that the inhibition of cellular transcription and translation, which rapidly develops in the SINV-infected cells, plays a critical role in preventing SG assembly. We mimicked virus-induced transcriptional shutoff by ActD treatment, and in agreement with the previously published data (46), within 2-4 h, it blocked SG formation in response to NaAs. The inhibition of SG assembly also requires the presence of nuclei and does not occur in enucleated cells. ActD-induced global transcriptional inhibition has been shown to cause the re-localization of multiple RNA-interacting proteins from the nuclei to the cytoplasm (46–49). This binding of RNA-interacting proteins to translationally inactive mRNAs is thought to render these RNA-protein complexes incapable of mediating SG formation (46). The transcriptional shutoff induced by SINV and CHIKV infections leads to the degradation of the entire pool of RPB1, the catalytic subunit of RNA polymerase II, and the relocalization of nuclear RNA-binding proteins to the cytoplasm (50). The accumulation of these proteins in the cytoplasm correlates with that found in ActD-treated cells. This relocalization of nuclear proteins to the cytoplasm provides a plausible explanation for the development of resistance to SG formation in the cells infected with OW alphaviruses. The ActD-induced inhibition of transcription also leads to the disintegration of preformed SGs (46), and thus, virus-induced transcriptional shutoff may explain the previously described dissolution of SG-like structures starting at 4 h p.i. with wt SFV (27). Mutant SINV variants, such as SINV/G/GFP and SINV/nsP2-683S/GFP, and the SINrep/L/GFP/Pac replicon, which exhibit the lower ability to inhibit cellular transcription, fail to block SG formation in response to NaAs.

The finding that translational shutoff is involved in the inhibition of SG development was unexpected. The cellular translation is inhibited within a few hours post infection with SINV or SFV (51, 52). This inhibition is mediated by two mechanisms. One of them depends on PKR and another is PKR-independent (21). The SG RNAs specific to SINV and SFV contain translational enhancers and remain efficiently translated in these conditions (53). The negative effect of virus-induced translation inhibition on the ability of cells to induce SG formation may be explained by the rapid reduction in the levels of one or more SG components or by a decrease in the rates of required posttranslational modifications. However, further investigation is needed to evaluate these hypotheses.

To date, there is no compelling evidence to suggest that SINV and CHIKV have developed specific mechanisms to interfere with SG formation. Instead, the virus-induced inhibitions of cellular transcription and translation play critical roles not only in downregulating the antiviral response but also in preventing SG development. In summary, the results of this new study demonstrate that: (i) the inhibition of SG formation during replication of SINV and likely other alphaviruses involves multiple mechanisms, (ii) the binding of G3BPs to CHIKV and SINV nsP3 HVDs, as well as the ADP-ribosylhydrolase activity of the nsP3 macro domain, are not critical contributors to the inhibition of SG development, at least when nsP3 is expressed at biologically relevant levels, (iii) the rapid transcriptional and translational shutoffs that occur in alphavirus-infected cells play major roles in inhibiting SG formation, and (iv) additional virus-specific mechanisms may also be involved in the suppression of SG formation, although this requires further investigation.

## Materials and Methods

### Cell cultures

NIH 3T3 cells were obtained from the American Type Culture Collection (Manassas, VA), and BHK-21 cells were provided by Paul Olivo (Washington University, St. Louis, MO). The NIH 3T3 *G3bp* dKO cell line was described elsewhere (20). All cell lines were maintained in alpha minimum essential medium (αMEM) supplemented with 10% fetal bovine serum (FBS) and vitamins at 37°C.

### Plasmid constructs

The original plasmids containing the infectious cDNAs of SINV/nsP3-GFP, SINV/GFP, SINV/G/GFP, SINV/nsP3-Cherry and CHIKV/GFP, as well as cDNAs of SINrep/L/GFP/Pac and SINrep/GFP replicons, and SINV helper genome were described elsewhere (15, 20, 35, 38, 54, 55). Modifications of nsP3 HVD in these plasmids and the expression constructs of nsP3 and HVD fusions in PiggyBac-based vector (20) were designed using standard PCR-based and cloning techniques. Details of the introduced modifications can be found in the Results section. VEEV replicons expressing GFP fused with SINV nsP3 or SINV HVD were designed based on a previously described VEEV replicon (56), using similar techniques. A VEEV helper-encoding plasmid was described elsewhere (57). Sequences of the plasmids and details of the cloning procedures are available upon request.

### Rescuing of recombinant viruses

Plasmids containing cDNAs of complete viral genomes, replicons, and helper genomes were purified by ultracentrifugation in CsCl gradients. The plasmids were linearized using unique restriction sites located downstream of the poly(A) tails. *In vitro* transcription was using SP6 RNA polymerase in the presence of a cap analog according to the manufacturer’s recommendations (New England Biolabs). The yields and integrities of the transcribed RNAs were analyzed by agarose gel electrophoresis under non-denaturing conditions. The transcription mixtures were used for transfection without further RNA purification. Viruses were rescued by electroporation of 3 µg of *in vitro*-synthesized RNAs into BHK-21 cells (58, 59) and harvested at 24 h post electroporation. In some experiments, viral replication was assessed immediately after RNA electroporation by seeding one-tenth of the electroporated cells into a 6-well Costar plate and harvesting media at the times indicated in the figures. Viral titers were determined using a standard plaque assay on BHK-21 cells (60).

RNA infectivities were assessed in the ICA. In this assay, ten-fold dilutions of electroporated BHK-21 cells were seeded into 6-well Costar plates containing subconfluent, naïve BHK-21 cells. After 2 h of incubation at 37°C, the media were replaced with 2 ml of MEM supplemented with 0.5% agarose and 3% FBS. Plaques were stained with crystal violet after 48 h of incubation at 37°C, and infectivity was determined in PFU per μg of transfected RNA.

### Packaging of alphavirus replicons

Replicons were packaged by co-electroporation of the *in vitro*-synthesized replicon and helper RNAs into BHK-21 cells, followed by incubation at 37°C (55). Infectious virions with replicons genomes were harvested at 24 h post electroporation.

Titers were determined by infecting BHK-21 cells in 6-well Costar plates (5×10^5^ cells/well) with serial dilutions of the harvested stocks in PBS supplemented with 1% FBS for 1 h. The inoculums were then replaced with complete BHK-21 media. After incubation at 37°C for 8 or 20 h, the numbers of infected, GFP-positive cells were evaluated under an inverted fluorescence Eclipse Ti Nikon microscope and used to calculate the titers in infectious units per ml (inf.u/ml).

### Analysis of viral replication

Equal numbers of cells of NIH 3T3 cells and their *G3bp* dKO derivative were seeded into 6-well Costar plates (5×10^5^ cells/well). After incubation for 4 h at 37°C, they were infected with the viruses at the MOIs described in the figure legends. At the indicated time points, the media were replaced, and viral titers were determined by plaque assay on BHK-21 cells.

To compare the abilities of the viruses to form plaques on NIH 3T3 and *G3bp* dKO cells, the latter cells were seeded into 6-well Costar plates (5×10^5^ cells/well). They were incubated at 37°C for 4 h, and then used for standard plaque assay. Plaques were stained with crystal violet after 48 h of incubation at 37°C.

### Stable cell lines

NIH 3T3 cells in 6-well Costar plates (3×10^5^ cells/well) were co-transfected with plasmids encoding GFP-nsP3 fusions and the integrase-encoding plasmid using Lipofectamine™ 3000 according to the manufacturer’s recommendations (ThermoFisher Scientific). Clones of blasticidin-resistant, GFP-positive cells were selected to express low levels of GFP fusions, which are more biologically relevant. The expression levels were further evaluated by WB and compared with the level of nsP3-GFP produced by SINV/nsP3-GFP recombinant virus in NIH 3T3 cells at 8 h p.i.

Cell line containing persistently replicating, noncytopathic SINrep/L/GFP/Pac replicon was generated by electroporating BHK-21 cells with the *in vitro*-synthesized replicon RNA. At 16 h post electroporation, the medium was supplemented with puromycin (5 μg/ml), and the cells were allowed to grow for 4 days before further experiments.

### Immunoprecipitations

NIH 3T3 cells in 6-well Costar plates were infected with packaged VEEV replicons at an MOI of 20 inf.u/cell. After incubation for 3 h in complete media, the cells were harvested, and protein complexes were isolated using anti-Flag MAb magnetic beads as previously described (61). Their compositions were analyzed by WB using the following antibodies: anti-Flag antibodies (F1804, Sigma) and anti-G3BP1 (gift from Dr. Richard Lloyd). Secondary antibodies labeled with Alexa Fluor™ Plus 680 or Alexa Fluor™ Plus 800 infrared dyes were acquired from ThermoFisher Scientific. Membranes were imaged on Odyssey Imaging System (LI-COR Biosciences).

### Western blotting

Proteins were separated using NuPAGE gels (ThermoFisher Scientific) and transferred to nitrocellulose membranes. After blocking in PBS supplemented with 5% nonfat milk or BSA, the membranes were incubated with antibodies specific to p-eIF2α (3398, Cell Signaling Technology), eIF2α (TA501313, OriGene), nsP2 (Mab7-4, custom), nsP3 (custom), β-actin (66009-1-Ig; Proteintech) and GFP (600-145-215, Rockland), followed by corresponding secondary antibodies labeled with infrared dyes. Membranes were imaged on Odyssey Imaging System (LI-COR Biosciences) and analyzed in Empiria Studio (LI-COR Biosciences).

### SG induction and immunofluorescence

Cells were seeded onto 8-well µ-slides (ibidi USA, Inc.), then infected at an MOI of 10 PFU/cell and incubated at 37°C for the times indicated in the figure legends. The presence of SGs was analyzed either without additional treatment or after induction of SG formation. To induce SGs, cells were treated with 0.75 mM NaAs for 45 min, and then fixed in 4% paraformaldehyde (PFA) in PBS for 20 min at room temperature. Cells were permeabilized with 0.5% Triton X-100 in PBS, blocked with 5 % BSA, and stained with antibodies specific to G3BP1 (gift from Dr. Richard Lloyd), eIF3b (sc-16377, Santa Cruz Biotechnology, Inc.) and TIAR (8509, Cell Signaling Technology), and corresponding fluorescent secondary antibodies. Images were acquired on a Zeiss LSM800 confocal microscope with a 63X 1.4NA PlanApochromat oil objective.

To analyze the effect of ActD-induced transcriptional shutoff on the ability of cells to generate SGs, NIH 3T3 cells in 8-well µ-slides (ibidi USA, Inc.), were incubated in complete media supplemented with ActD (5 µg/ml) for 1, 2, and 4 h. Then, they were treated with NaAs and stained with TIAR– and eIF3b-specific antibodies as described above.

## ACKNOWLEDGMENTS

We thank Nikita Shiliaev for technical assistance. This study was supported by Public Health Service grants R01AI133159 and R01AI118867 to E.I.F. as well as R01AI050537, R21AI146969 and R01AI073301 to I.F. and by the UAB Research Acceleration Funds to E.I.F and I.F.

## REFERENCES

1. Strauss JH, Strauss EG. 1994. The alphaviruses: gene expression, replication, evolution. Microbiol Rev 58:491–562.

2. Weaver SC, Barrett AD. 2004. Transmission cycles, host range, evolution and emergence of arboviral disease. Nat Rev Microbiol 2:789–801.

3. Weaver SC, Frolov I. 2005. Togaviruses, p 1010–1024. In Mahy BWJ, Meulen Vt (ed), Virology, vol 2. ASM Press, Salisbury, UK.

4. Weaver SC, Lecuit M. 2015. Chikungunya Virus Infections. N Engl J Med 373:94–5.

5. Lemm JA, Bergqvist A, Read CM, Rice CM. 1998. Template-dependent initiation of Sindbis virus replication in vitro. J Virol 72:6546–6553.

6. Lemm JA, Rice CM. 1993. Assembly of functional Sindbis virus RNA replication complexes: Requirement for coexpression of P123 and P34. J Virol 67:1905–1915.

7. Lemm JA, Rice CM. 1993. Roles of nonstructural polyproteins and cleavage products in regulating Sindbis virus RNA replication and transcription. J Virol 67:1916–1926.

8. Shirako Y, Strauss JH. 1994. Regulation of Sindbis virus RNA replication: Uncleaved P123 and nsP4 function in minus strand RNA synthesis whereas cleaved products from P123 are required for efficient plus strand RNA synthesis. J Virol 185:1874–1885.

9. Rice CM, Strauss JH. 1981. Nucleotide sequence of the 26S mRNA of Sindbis virus and deduced sequence of the encoded virus structural proteins. Proc Natl Acad Sci USA 78:2062–2066.

10. Strauss EG, Rice CM, Strauss JH. 1984. Complete nucleotide sequence of the genomic RNA of Sindbis virus. Virology 133:92–110.

11. Garmashova N, Gorchakov R, Volkova E, Paessler S, Frolova E, Frolov I. 2007. The Old World and New World alphaviruses use different virus-specific proteins for induction of transcriptional shutoff. J Virol 81:2472–84.

12. Atasheva S, Garmashova N, Frolov I, Frolova E. 2008. Venezuelan equine encephalitis virus capsid protein inhibits nuclear import in Mammalian but not in mosquito cells. J Virol 82:4028–41.

13. Akhrymuk I, Lukash T, Frolov I, Frolova EI. 2019. Novel Mutations in nsP2 Abolish Chikungunya Virus-Induced Transcriptional Shutoff and Make the Virus Less Cytopathic without Affecting Its Replication Rates. J Virol 93.

14. Akhrymuk I, Frolov I, Frolova EI. 2018. Sindbis Virus Infection Causes Cell Death by nsP2-Induced Transcriptional Shutoff or by nsP3-Dependent Translational Shutoff. J Virol 92.

15. Frolova EI, Fayzulin RZ, Cook SH, Griffin DE, Rice CM, Frolov I. 2002. Roles of nonstructural protein nsP2 and Alpha/Beta interferons in determining the outcome of Sindbis virus infection. J Virol 76:11254–64.

16. Frolov I, Schlesinger S. 1994. Comparison of the effects of Sindbis virus and Sindbis virus replicons on host cell protein synthesis and cytopathogenicity in BHK cells. J Virol 68:1721–1727.

17. Gorchakov R, Frolova E, Frolov I. 2005. Inhibition of transcription and translation in Sindbis virus-infected cells. J Virol 79:9397–409.

18. Gorchakov R, Garmashova N, Frolova E, Frolov I. 2008. Different types of nsP3-containing protein complexes in Sindbis virus-infected cells. J Virol 82:10088–101.

19. Frolova EI, Gorchakov R, Pereboeva L, Atasheva S, Frolov I. 2010. Functional Sindbis virus replicative complexes are formed at the plasma membrane. J Virol 84:11679–95.

20. Kim DY, Reynaud JM, Rasalouskaya A, Akhrymuk I, Mobley JA, Frolov I, Frolova EI. 2016. New World and Old World Alphaviruses Have Evolved to Exploit Different Components of Stress Granules, FXR and G3BP Proteins, for Assembly of Viral Replication Complexes. PLoS Pathog 12:e1005810.

21. Gorchakov R, Frolova E, Williams BR, Rice CM, Frolov I. 2004. PKR-dependent and – independent mechanisms are involved in translational shutoff during Sindbis virus infection. J Virol 78:8455–67.

22. Akhrymuk I, Frolov I, Frolova EI. 2016. Both RIG-I and MDA5 detect alphavirus replication in concentration-dependent mode. Virology 487:230–41.

23. Kedersha N, Ivanov P, Anderson P. 2013. Stress granules and cell signaling: more than just a passing phase? Trends Biochem Sci 38:494–506.

24. Kedersha N, Chen S, Gilks N, Li W, Miller IJ, Stahl J, Anderson P. 2002. Evidence that ternary complex (eIF2-GTP-tRNA(i)(Met))-deficient preinitiation complexes are core constituents of mammalian stress granules. Mol Biol Cell 13:195–210.

25. Kedersha NL, Gupta M, Li W, Miller I, Anderson P. 1999. RNA-binding proteins TIA-1 and TIAR link the phosphorylation of eIF-2 alpha to the assembly of mammalian stress granules. J Cell Biol 147:1431–42.

26. Poblete-Duran N, Prades-Perez Y, Vera-Otarola J, Soto-Rifo R, Valiente-Echeverria F. 2016. Who Regulates Whom? An Overview of RNA Granules and Viral Infections. Viruses 8.

27. McInerney GM, Kedersha NL, Kaufman RJ, Anderson P, Liljestrom P. 2005. Importance of eIF2alpha phosphorylation and stress granule assembly in alphavirus translation regulation. Mol Biol Cell 16:3753–63.

28. Panas MD, Varjak M, Lulla A, Eng KE, Merits A, Karlsson Hedestam GB, McInerney GM. 2012. Sequestration of G3BP coupled with efficient translation inhibits stress granules in Semliki Forest virus infection. Mol Biol Cell 23:4701–12.

29. Jayabalan AK, Adivarahan S, Koppula A, Abraham R, Batish M, Zenklusen D, Griffin DE, Leung AKL. 2021. Stress granule formation, disassembly, and composition are regulated by alphavirus ADP-ribosylhydrolase activity. Proc Natl Acad Sci U S A 118.

30. Tossavainen H, Aitio O, Hellman M, Saksela K, Permi P. 2016. Structural Basis of the High Affinity Interaction between the Alphavirus Nonstructural Protein-3 (nsP3) and the SH3 Domain of Amphiphysin-2. J Biol Chem 291:16307–17.

31. Panas MD, Ahola T, McInerney GM. 2014. The C-terminal repeat domains of nsP3 from the Old World alphaviruses bind directly to G3BP. J Virol 88:5888–93.

32. Fros JJ, Domeradzka NE, Baggen J, Geertsema C, Flipse J, Vlak JM, Pijlman GP. 2012. Chikungunya virus nsP3 blocks stress granule assembly by recruitment of G3BP into cytoplasmic foci. J Virol 86:10873–9.

33. Meshram CD, Agback P, Shiliaev N, Urakova N, Mobley JA, Agback T, Frolova EI, Frolov I. 2018. Multiple Host Factors Interact with the Hypervariable Domain of Chikungunya Virus nsP3 and Determine Viral Replication in Cell-Specific Mode. J Virol 92.

34. McInerney GM. 2015. FGDF motif regulation of stress granule formation. DNA Cell Biol 34:557–60.

35. Frolova E, Gorchakov R, Garmashova N, Atasheva S, Vergara LA, Frolov I. 2006. Formation of nsP3-specific protein complexes during Sindbis virus replication. J Virol 80:4122–34.

36. Akhrymuk I, Frolov I, Frolova EI. 2018. Sindbis Virus Infection Causes Cell Death by nsP2-Induced Transcriptional Shutoff or by nsP3-Dependent Translational Shutoff. J Virol 92:e01388.

37. Frolov I, Agapov E, Hoffman TA, Jr., Pragai BM, Lippa M, Schlesinger S, Rice CM. 1999. Selection of RNA replicons capable of persistent noncytopathic replication in mammalian cells. J Virol 73:3854–65.

38. Frolov I, Agapov E, Hoffman Jr. TA, Prágai BM, Lippa M, Schlesinger S, Rice CM. 1999. Selection of RNA replicons capable of persistent noncytopathic replication in mammalian cells. J Virol 73:3854–3865.

39. Frolov I, Schlesinger S. 1994. Translation of Sindbis virus mRNA: effects of sequences downstream of the initiating codon. J Virol 68:8111–7.

40. Nikonov A, Molder T, Sikut R, Kiiver K, Mannik A, Toots U, Lulla A, Lulla V, Utt A, Merits A, Ustav M. 2013. RIG-I and MDA-5 detection of viral RNA-dependent RNA polymerase activity restricts positive-strand RNA virus replication. PLoS Pathog 9:e1003610.

41. Froshauer S, Kartenbeck J, Helenius A. 1988. Alphavirus RNA replicase is located on the cytoplasmic surface of endosomes and lysosomes. J Cell Biol 107:2075–86.

42. Frolov I, Kim DY, Akhrymuk M, Mobley JA, Frolova EI. 2017. Hypervariable Domain of Eastern Equine Encephalitis Virus nsP3 Redundantly Utilizes Multiple Cellular Proteins for Replication Complex Assembly. J Virol 91.

43. Li C, Debing Y, Jankevicius G, Neyts J, Ahel I, Coutard B, Canard B. 2016. Viral Macro Domains Reverse Protein ADP-Ribosylation. J Virol 90:8478–86.

44. Abraham R, Hauer D, McPherson RL, Utt A, Kirby IT, Cohen MS, Merits A, Leung AKL, Griffin DE. 2018. ADP-ribosyl-binding and hydrolase activities of the alphavirus nsP3 macrodomain are critical for initiation of virus replication. Proc Natl Acad Sci U S A 115:E10457–E10466.

45. Eckei L, Krieg S, Butepage M, Lehmann A, Gross A, Lippok B, Grimm AR, Kummerer BM, Rossetti G, Luscher B, Verheugd P. 2017. The conserved macrodomains of the non-structural proteins of Chikungunya virus and other pathogenic positive strand RNA viruses function as mono-ADP-ribosylhydrolases. Sci Rep 7:41746.

46. Bounedjah O, Desforges B, Wu TD, Pioche-Durieu C, Marco S, Hamon L, Curmi PA, Guerquin-Kern JL, Pietrement O, Pastre D. 2014. Free mRNA in excess upon polysome dissociation is a scaffold for protein multimerization to form stress granules. Nucleic Acids Res 42:8678–91.

47. Fan XC, Steitz JA. 1998. HNS, a nuclear-cytoplasmic shuttling sequence in HuR. Proc Natl Acad Sci U S A 95:15293–8.

48. Zhang T, Delestienne N, Huez G, Kruys V, Gueydan C. 2005. Identification of the sequence determinants mediating the nucleo-cytoplasmic shuttling of TIAR and TIA-1 RNA-binding proteins. J Cell Sci 118:5453–63.

49. Caceres JF, Screaton GR, Krainer AR. 1998. A specific subset of SR proteins shuttles continuously between the nucleus and the cytoplasm. Genes Dev 12:55–66.

50. Akhrymuk I, Kulemzin SV, Frolova EI. 2012. Evasion of the innate immune response: the Old World alphavirus nsP2 protein induces rapid degradation of Rpb1, a catalytic subunit of RNA polymerase II. J Virol 86:7180–91.

51. McInerney GM, Smit JM, Liljestrom P, Wilschut J. 2004. Semliki Forest virus produced in the absence of the 6K protein has an altered spike structure as revealed by decreased membrane fusion capacity. Virology 325:200–6.

52. Frolov I, Schlesinger S. 1994. Comparison of the effects of Sindbis virus and Sindbis virus replicons on host cell protein synthesis and cytopathogenicity in BHK cells. J Virol 68:1721–7.

53. Frolov I, Schlesinger S. 1996. Translation of Sindbis virus mRNA: analysis of sequences downstream of the initiating AUG codon that enhance translation. J Virol 70:1182–90.

54. Foy NJ, Akhrymuk M, Akhrymuk I, Atasheva S, Bopda-Waffo A, Frolov I, Frolova EI. 2013. Hypervariable domains of nsP3 proteins of New World and Old World alphaviruses mediate formation of distinct, virus-specific protein complexes. J Virol 87:1997–2010.

55. Bredenbeek PJ, Frolov I, Rice CM, Schlesinger S. 1993. Sindbis virus expression vectors: packaging of RNA replicons by using defective helper RNAs. J Virol 67:6439–46.

56. Petrakova O, Volkova E, Gorchakov R, Paessler S, Kinney RM, Frolov I. 2005. Noncytopathic replication of Venezuelan equine encephalitis virus and eastern equine encephalitis virus replicons in Mammalian cells. J Virol 79:7597–608.

57. Volkova E, Gorchakov R, Frolov I. 2006. The efficient packaging of Venezuelan equine encephalitis virus-specific RNAs into viral particles is determined by nsP1-3 synthesis. Virology 344:315–27.

58. Gorchakov R, Hardy R, Rice CM, Frolov I. 2004. Selection of functional 5’ cis-acting elements promoting efficient sindbis virus genome replication. J Virol 78:61–75.

59. Liljestrom P, Garoff H. 1991. A new generation of animal cell expression vectors based on the Semliki Forest virus replicon. Biotechnology (N Y) 9:1356–61.

60. Atasheva S, Krendelchtchikova V, Liopo A, Frolova E, Frolov I. 2010. Interplay of acute and persistent infections caused by Venezuelan equine encephalitis virus encoding mutated capsid protein. J Virol 84:10004–15.

61. Dominguez F, Shiliaev N, Lukash T, Agback P, Palchevska O, Gould JR, Meshram CD, Prevelige PE, Green TJ, Agback T, Frolova EI, Frolov I. 2021. NAP1L1 and NAP1L4 Binding to Hypervariable Domain of Chikungunya Virus nsP3 Protein Is Bivalent and Requires Phosphorylation. J Virol 95:e0083621.

